# Cholinergic Signaling Modulates Intestinal Pathophysiology in a *Drosophila* Model of Cystic Fibrosis

**DOI:** 10.1101/2025.07.02.662792

**Authors:** Elizabeth A. Lane, Afroditi Petsakou, Ying Liu, Weihang Chen, Mujeeb Qadiri, Yanhui Hu, Norbert Perrimon

## Abstract

Cystic fibrosis (CF) is a monogenic genetic disease caused by mutations in the Cystic Fibrosis Transmembrane conductance Regulator (CFTR) chloride/bicarbonate channel, which is expressed in certain epithelial cells. Current therapies focus on restoring CFTR function, but many gut-related pathologies persist, highlighting the need for complementary treatments to improve the quality of life of people with CF. In this study, we use *Drosophila melanogaster* as a model to investigate the gut-specific effects of *Cftr* loss. We demonstrate that enterocyte specific knockdown of *Cftr* in flies recapitulates several CF pathologies, including reduced intestinal motility, nutrient malabsorption, and decreased energy stores. Using single-nuclei RNA sequencing (snRNA-seq), we identify significant transcriptional changes in the CF model gut, including the upregulation of *acetylcholine esterase* (*Ace,* human *AChE*), which leads to reduced cholinergic signaling. Cholinergic signaling has been shown to affect CFTR function but this is the first time CFTR loss of function has been shown to alter cholinergic signaling. Functional assays confirm that cholinergic sensitivity is diminished in CF guts. Furthermore, restoring cholinergic signaling via *Ace* knockdown rescues multiple CF-associated phenotypes. Additionally, we identify the transcription factor Fork head (Fkh), the *Drosophila* homolog of human FOXA1/FOXA2, which is known to be a positive regulator of *Cftr* transcription in the intestine, as a positive regulator of *Ace* expression in CF guts. This study establishes the *Drosophila* gut as a powerful model to investigate CF pathogenesis, genetic modifiers, and identifies *Ace* and *fkh* as genetic modifiers. This work also suggests that enhancing cholinergic signaling may represent a viable therapeutic strategy for gastrointestinal manifestations of CF.

**Author Summary:** Cystic fibrosis (CF) is a genetic disease that causes complications in multiple organ systems, including the lungs and gastrointestinal tract. While recent therapies have greatly improved life span and respiratory outcomes in people with CF (pwCF), they continue to report significant gastrointestinal symptoms that impact their quality of life. Therefore, there is a need to better understand the progression of CF in the gut and identify gut specific therapeutic targets to improve patient quality of life. In this study we use *Drosophila* to model gut specific complications of CF. We show our model recapitulates many CF clinical presentations in the gut demonstrating our model may be clinically relevant. Furthermore, we identify *acetylcholine esterase* (*Ace*) as a gene that is increased in our CF model guts that is important for the development of CF pathologies. We additionally, perform a screen to identify a transcriptional regulator of *Ace*, whose genetic manipulation also regulates CF phenotypes. Our findings not only identify a potential target to alleviate GI symptoms in patients with CF, but also validate the *Drosophila* gut as a robust model for studying CF pathogenesis, genetic modifiers, and screening therapeutics.

## Introduction

Cystic fibrosis (CF) is a genetic disease that affects approximately 1 in 2,500 newborns in the United States(1). CF is caused by mutations in the Cystic Fibrosis Transmembrane conductance Regulator (CFTR) chloride channel, which is expressed in certain epithelial tissues, including the respiratory tract, pancreas, and the gut(2–4). The CF phenotype is most closely associated with the accumulation of mucus in the pulmonary and gastrointestinal tracts, which can lead to inflammation, bacterial infection, and malnutrition(1, 5–8). Much of CF research has focused on the lung, as the primary cause of death in people with CF (pwCF) have been related to lung complications, however, CF causes pathologies in other CFTR-expressing tissues, including the gut, that result in various clinical phenotypes(7, 9–13). For example, many pwCF are born with intestinal blockage (Meconium Ileus) and later in life have decreased intestinal motility, dysbiosis, inflammation, and poor nutrient absorption in the gut(7, 9–13). Additionally, as life expectancy has increased following better treatments, complications in the gastrointestinal tract have become an increasing cause of morbidity(10–15). These complications include small bowel bacterial overgrowth, bowel obstructions, and increased risk of intestinal cancers(11–13). Recent studies have suggested that CF modulator drugs may not fully correct the inflammation and dysbiosis seen in the gut of pwCF and that pwCF on modulator drugs still report GI symptoms(14, 16–19). Therefore, additional therapies targeting the gut, not directly related to modulating CF activity may be useful for quality of life of pwCF. Recently, there have been studies highlighting how the CF gut can crosstalk with many other organ systems, further highlighting the importance of investigating how loss of CFTR function effects gut biology(20–22).

The *Drosophila* midgut, analogous to the human intestine, is comprised primarily of 4 main cell types: the absorptive enterocytes (ECs), the neuropeptide producing enteroendocrine (EE) cells, intestinal stem cells (ISCs), and the intermediate progenitor enteroblasts (EBs) (23). The *Drosophila* midgut has been used to study a number of questions in stem cell biology, cell signaling, and physiology, and many of the mechanisms active in the intestines of flies have already been shown to apply more broadly to other organisms and may therefore be relevant for human pathologies(23–25). The fly ortholog of human CFTR has recently been identified and used to establish an intestinal CF model(26). This model revealed that *Cftr* knockdown in the *Drosophila* intestine disrupts osmotic homeostasis and displays CF-like phenotypes in the intestinal epithelium(26). While this work introduced *Drosophila* as a CF model, the phenotypes examined were largely cellular phenotypes, such as increased chloride and sodium levels in the ECs, and not clinical manifestations of CF.

Here, we further characterize the CF model gut, demonstrating that several clinical pathologies, including reduced intestinal motility and nutrient malabsorption are preserved. Furthermore, the *Drosophila* CF model displays characteristics consistent with a failure to thrive phenotype. Additionally, we perform single nuclei RNA-seq (snRNA-seq) to further characterize the CF gut model and learn new biology relevant to CF. Interestingly, we found increased *acetylcholine esterase* (*Ace*) expression in CF model guts, which results in reduced cholinergic signaling potential. Recent work in *Drosophila* has demonstrated that cholinergic signaling is important for maintenance of the intestinal epithelial barrier and is required for the intestinal epithelium to return to homeostasis after injury(27, 28). Additionally cholinergic signaling has been implicated in other diseases of intestinal inflammation such as Inflammatory Bowel Disease (IBD) but has not yet been investigated in the context of the CF intestine(29, 30). Previous studies have demonstrated that cholinergic signaling can increase CFTR function(31–34) but this is the first work that demonstrates CFTR function can have a reciprocal effect on cholinergic signaling. We further show that the decrease in cholinergic signaling observed in the CF model guts may be clinically relevant as restoring sensitivity to cholinergic signaling rescues many CF pathologies in *Drosophila*. Finally, we identify Fork head (Fkh), a FOXA1/A2 homolog as a transcriptional regulator of *Ace* expression in the CF model gut. FOXA1/A2 is a pioneering transcription factor that positively regulates the expression of *Cftr* and other transmembrane proteins and ion channels important for regulating ion homeostasis in the intestinal epithelium(35, 36). Altogether, our findings help establish *Drosophila* as a model organism to study GI manifestations of CF. We identify Ace (human: AChE) and Fkh (human: FOXA1/A2) as genetic modifiers of CF and our findings suggests that the cholinergic signaling pathway may be a viable therapeutic target in CF gastrointestinal disease.

## Results

### Knockdown of *Cftr* in the *Drosophila* midgut recapitulates clinical pathologies of Cystic Fibrosis

People with CF (pwCF) display a range of multiple organ specific as well as systemic pathologies, including gastrointestinal complications(10–13, 37–39). Therefore, we examined whether *Drosophila* CF model guts, with enterocyte-specific knockdown of *Cftr* (*Myo1A-Gal4 > UAS-Cftr^RNAi^*) recapitulated typical clinical pathologies. pwCF are at increased risk for gastrointestinal cancers and microbial dysbiosis(11, 13, 39, 40). These clinical observations are consistent with previous findings in the *Drosophila* CF model, showing gut hyperplasia and increased bacterial load(26). Additional common CF gut pathologies such as meconium ileus, distal intestinal obstruction syndrome, and constipation are linked to impaired intestinal motility(9, 38, 41). To test whether the CF model exhibits reduced gut motility, we performed an excretion assay. *Cftr* knockdown flies showed significantly reduced excretion compared to wild-type (WT) controls, indicating impaired intestinal transit (Fig 1A, S1A). Malabsorption of nutrients in the intestine is another hallmark of CF(42). In line with this, CF model flies exhibited increased glucose, triacyl glycerides (TAGs), and protein levels in their excreta compared to WT flies, suggesting reduced nutrient absorption in the intestine (Fig 1B-D, S1B). Of note increased tissue breakdown and stress responses can result in the fly kidney (Malpighian tubules) depositing lipids into the excreta which may contribute to the increased nutrients we observe in the excreta in our model(43).

**Figure 1.**
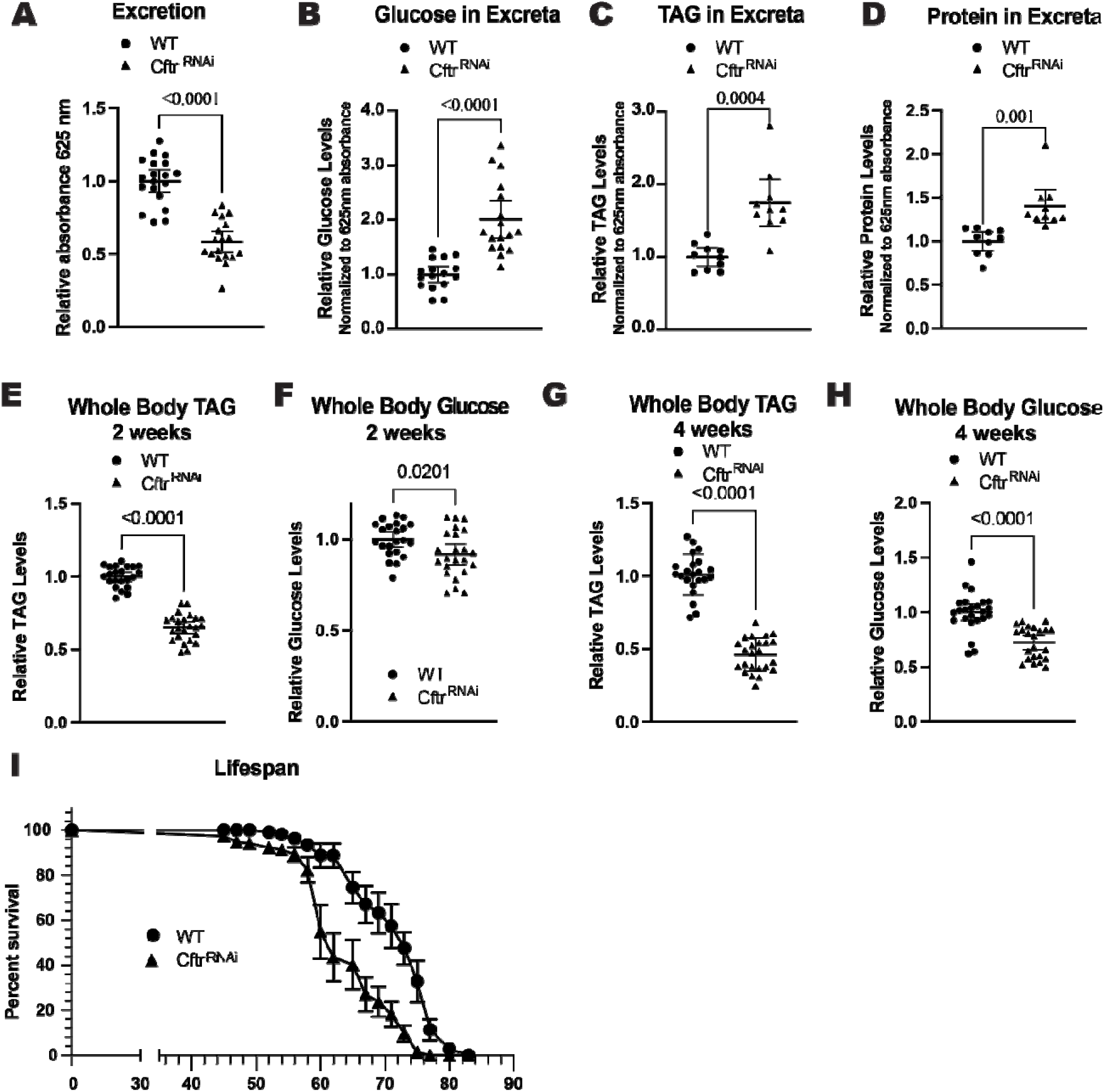
CF model gut recapitulates hallmarks of CF. (**A**) CF model guts have decreased excretion rate compared to WT guts. n= 19 vials of 10-15 females from 3 independent experiments. (**B**) CF model guts have increased glucose in excreta compared to WT flies. n= 16 (WT), 17 (*Cftr^RNAi^*) vials of 15-20 females from 3 independent experiments. (**C-D**) CF model guts have increased TAG (**C**) and protein (**D**) in excreta compared to WT flies. n= 10 vials of 60 females from 3 independent experiments. (**E**) CF model guts have reduced TAG levels at 2 weeks of age compared to WT flies. n= 22 (WT), 24 (*Cftr^RN^*^Ai^) of 5 pooled females from 4 independent experiments. (**F**) CF model guts have slightly reduced whole body glucose levels at 2 weeks of age compared to WT flies. n= 22 (WT), 24 (*Cftr^RNAi^*) of 5 pooled females from 4 independent experiments. (**G**) CF model gut flies have reduced whole body TAG levels at 4 weeks of age compared to WT flies. n= 22 (WT), 24 (*Cftr^RNAi^*) of 5 pooled females from 4 independent crosses. (**H**) CF model gut flies have reduced whole body glucose at 4 weeks of age compared to WT flies. n= 24 (WT), 23 (*Cftr^RNAi^*) of 5 pooled females from 4 independent crosses. (**A-H**) pValues were calculated using the Mann-Whitney test in GraphPad prism. Error bars are mean with 95% CI. (**I**) CF model gut flies have reduced lifespans compared to WT flies. n= 9 (WT), 12 (*Cftr^RNAi^*) vials with 10-15 females from 2 independent crosses, error bars are mean +/- SEM. (**A-I**) WT is *Myo1A > +* and *Cftr^RNA^*^i^ is *Myo1A >UAS-Cftr^RNAi^*.

Failure to thrive, where pwCF fail to gain weight and grow at the expected rate, is a more systemic pathology with a strong intestinal component(7, 42, 44). *Cftr* knockdown flies have significantly reduced energy stores, with lower levels of whole-body TAG and glucose at both 2 and 4 weeks of age (Fig 1E-H, S1C-F). The reduction of TAG levels is robust at both 2 and 4 weeks of age while the reduction of glucose stores is more apparent at 4 weeks of age. This reduction in whole body energy stores is not due to a developmental defect as inducing *Cftr* knockdown in adult flies using the temperature sensitive Gal80^ts^ repressor also leads to reduced whole body TAGs and glucose (S1 G-H). Furthermore, the reduction in whole body energy stores is not due to reduced food intake as the CF gut model flies consume similar amounts of food to WT flies over 24 hrs and in 30 minutes after starvation (S1I-J). This decrease in whole body energy stores is consistent with a failure to thrive phenotype. As the CF gut model only has reduced *Cftr* expression in the midgut, it may be useful in studying gut-specific contributions to the failure to thrive outcome.

Finally, pwCF have reduced lifespans, although they have improved dramatically in recent years due to the availability of better treatments(45). Our model fly shows that loss of *Cftr* in the gut alone is sufficient to reduce the lifespan of the fly, highlighting the importance of CFTR function in the gut (Fig 1I, S1K). Overall, these results show that loss of *Cftr* in the enterocytes recapitulates many hallmarks of CF in the gastrointestinal system.

### Single nuclei analysis of CF model guts

To further characterize the CF gut model, we performed snRNA-seq on the midgut of WT and CF model gut flies. The sn-RNA-seq analysis recovered 3,669 nuclei for WT and 4,166 for CF midguts that were clustered into 21 clusters annotated using the marker genes from the previously published cell atlas of the *Drosophila* gut(46, 47) (Fig 2A). These clusters include multiple enterocyte (EC) clusters that map along the *Drosophila* digestive tract and show transcriptional similarity to mammalian enterocytes(48) (S2A). The posterior EC (pEC) clusters show high transcriptional similarity to the mature distal enterocytes while the anterior ECs (aEC) cluster only shows moderate similarity to mature proximal enterocytes (S2A). The middle ECs (mECs), copper cells, and large flat cells (LFC) which reside in the middle of the *Drosophila* midgut show moderate similarity to mammalian EC clusters (S2A). The *Drosophila* midgut does not contain specific goblet or paneth cells but the aEC clusters are transcriptionally similar to paneth cells and multiple EC clusters express genes specific to goblet cells in the mammalian gut (S2A). The *Drosophila* enteroendocrine cells show strong sequence similarity to the mammalian enteroendocrine cells. Finally, the *Drosophila*, intestinal stem cells (ISCs), enteroblasts (EBs), and adult differentiating enterocytes are similar to the mammalian stem cells, transit amplifying cells, and enterocyte progenitor cells (S2A).

**Figure 2.**
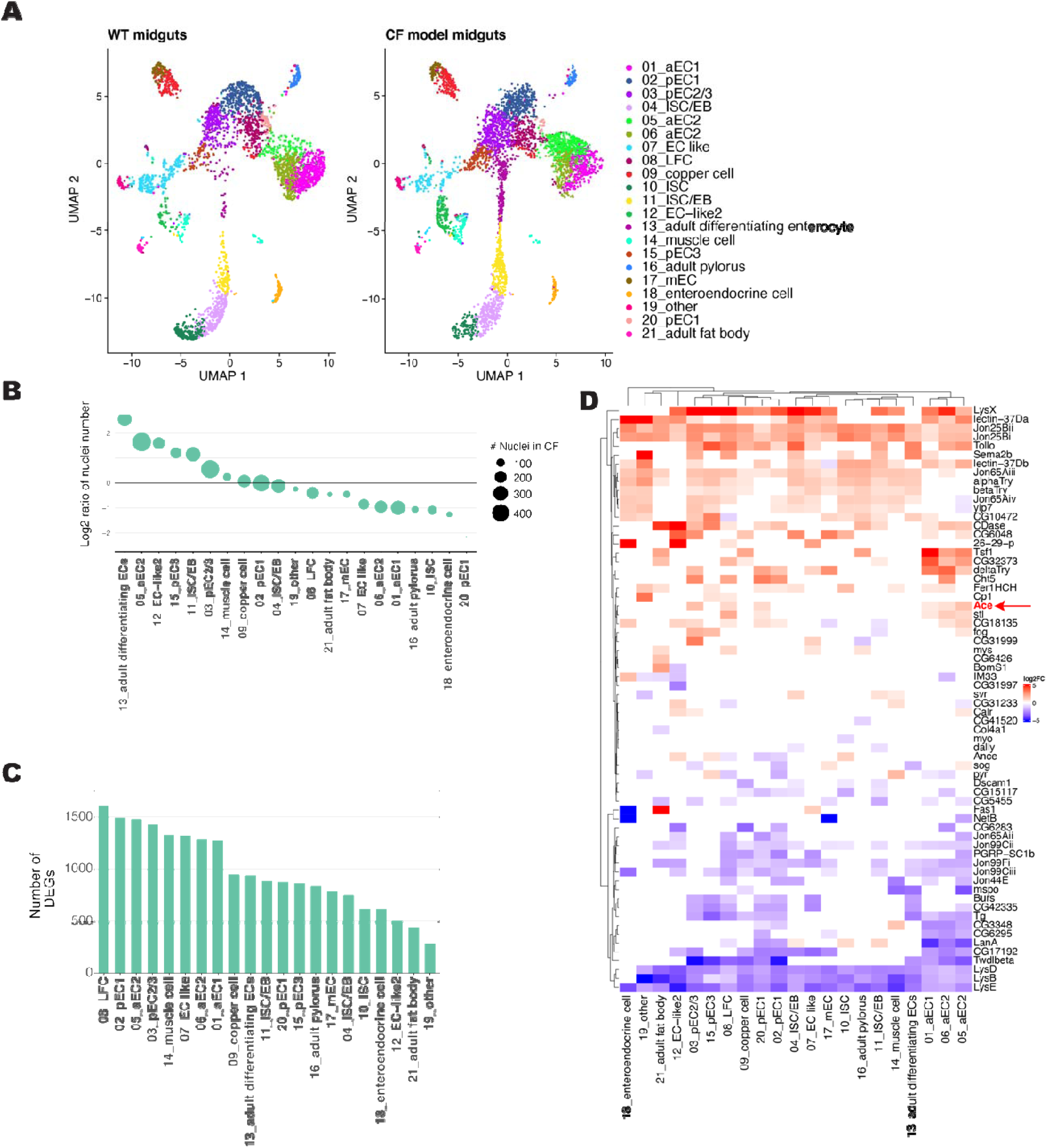
snRNA-Seq of CF model guts. **(A)** Annotated cell clusters of snRNA-Seq from WT (*Myo1A> +*) and CF (*Myo1A>Cftr^RNAi^*) midguts, visualized with UMAP. (**B**) Over and underrepresented cell clusters in CF model gut snRNA-seq data set. Size of nuclei represent number of nuclei in CF model gut analysis for indicated cluster. (**C**) Number of differentially expressed genes in each cell cluster between WT and CF model gut snRNA-seq data sets. (**D**) Heat map of the top 5 differentially expressed secreted proteins for each cell type. Color reflects the log2fold change in expression in CF model guts comparing to control per cell cluster.

UMAP analysis of the sn-RNA-seq data does not identify any unique clusters in the CF model guts, however there are a number of clusters that are over or under-represented in the CF model (Fig 2A-B, S2B). This includes increased ISCs, which is consistent with the hyperplasia previously described in the *Drosophila* CF model gut(26). Furthermore, there are a significant number of differentially expressed genes (DEGs) between WT and CFTR deficient guts across all identified cell clusters (Fig 2C, File S1). Together these results indicate substantial differences in the transcriptional profile and cellular composition between WT and CF model guts.

Interestingly, many secreted peptides are differentially expressed indicating that the CF gut model may be a good model to study the crosstalk between the CF gut and other organs (Fig 2D, S2C, File S2). Among the upregulated secreted proteins in the CF model midguts are several that have also been identified as key factors secreted by Yki-activated gut tumors involved in tumorigenesis and communication with other organs (*Impl2*, *Pvf1*, *Itp*, and *Upd3*)(49, 50) (S2D). This overlap suggests that the CF model gut may exhibit a predisposition toward intestinal tumorigenesis similar to pwCF who are at increased risk of gastrointestinal cancers(11, 13, 39, 40).

In addition to the expected increase of cells in the ISCs cell clusters(26) there is a redistribution of cells in anterior and posterior EC cell clusters (Fig 2A-B, S2B). In the CF preferred anterior EC cluster, one of the top 5 differentially expressed secreted peptides is Acetylcholine esterase (Ace, human AChE) (Fig 2D). *Ace* expression is increased overall in CF model guts but is particularly apparent in the anterior EC clusters (Fig 3A, S3A-B). We confirmed increased *Ace* expression in our CF model guts by RT-qPCR analysis of whole guts (S3C).

**Figure 3.**
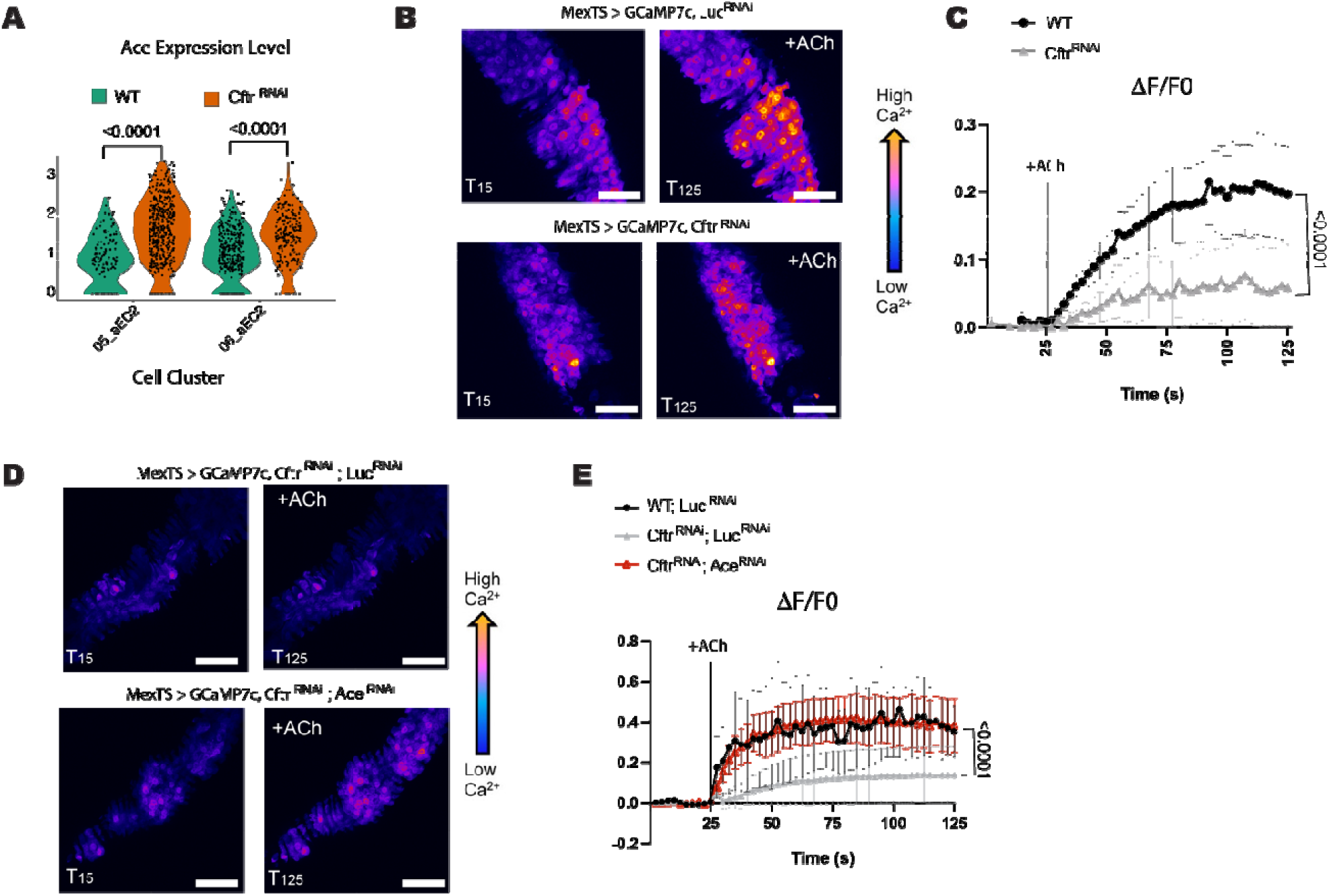
CF model guts have reduced sensitivity to cholinergic signaling. **(A)** Violin plot of acetylcholine esterase (*Ace*) expression in anterior EC2 cell clusters indicates higher *Ace* expression in CF model guts. Each dot represents expression in a single-nucleus, n=137 (WT) or 479 (*Cftr^RNAi^*) for 05_aEC2 and n=362 (WT) or 213 (*Cftr^RNAi^*) for 06_aEC2. pValues were calculated using the Mann-Whitney test in ggplot2. **(B-C)** *Cftr* deficient midguts have reduced sensitivity to acetylcholine (ACh) stimulation as measured by calcium levels in enterocytes. **(B)** Representative images of GCaMP7c fluorescence, in WT (*MexTS > GCaMP7c, LucRNAi*) and *Cftr* deficient guts (*MexTS > GCaMP7c, Cftr^RNAi^*) before (T15s) or after addition of ACh (T125). Scale bar 50 μm. **(C)** Graph of average relative fluorescent intensity, ΔF/F0, per frame (2.5s per frame) and genotype. n= 9 (WT) and 10 (*Cftr^RNAi^*) from 4 independent experiments. Error bars are mean +/- SEM. pValue was calculated using the Mann-Whitney test in GraphPad prism. **(D-E)** *Ace* knockdown increases sensitivity to ACh stimulation in *Cftr* deficient enterocytes. **(D)** Representative images of GCaMP7c fluorescence, in *Cftr* deficient (*MexTS > GCaMP7c, Cftr^RNAi^; Luc^RNAi^*) midguts and *Cftr* deficient guts with *Ace* knockdown (*MexTS > GCaMP7c, Cftr^RNAi^; Ace^RNAi^*) before (T15s) or after addition of ACh (T125). Scale bar 50 μm. **(E)** Graph of average relative fluorescent intensity, ΔF/F0, per frame (2.5s per frame) and genotype. n= 3 (WT), 5 (*Cftr^RNAi^*, *Luc^RNAi^*), 4 (*Cftr^RNAi^*, *Ace^RNAi^*) from 3 independent experiment. Error bars are mean +/- SEM. pValues were calculated using 2-way ANOVA with Tukey’s multiple comparisons main column effect in GraphPad prism.

Ace hydrolyzes acetylcholine into acetate and choline thereby inhibiting cholinergic signaling. Cholinergic signaling has recently been shown to be required for recovery of the intestinal epithelium after damage and for maintaining intestinal barrier function in *Drosophila*(27, 28). Furthermore, cholinergic signaling has been shown to increase CFTR activity but there are no studies showing CFTR function affecting cholinergic signaling(31–34). Therefore, we investigated the importance of increased *Ace* expression in our CF gut model.

### Reduced Cholinergic signaling in the CF Gut Model

As Ace degrades acetylcholine (ACh), the agonist of cholinergic receptors, increased *Ace* expression should decrease sensitivity to ACh and limit cholinergic signaling(27). Cholinergic signaling has been proposed to use Ca^2+^ as a secondary messenger to modulate the mammalian intestinal epithelium(51). Therefore, to test for cholinergic sensitivity in the *Cftr* deficient midguts we visualized Ca^2+^ by conditionally expressing the Ca^2+^ indicator GCaMP7c(52) in ECs and performed *ex vivo* live imaging(27). We found that Ca^2+^ levels in CF model guts were reduced in response to ACh, indicating that CF model guts had a dampened response to ACh stimulation compared to WT guts (Fig 3B-C).

Cholinergic receptors include both muscarinic and nicotinic receptors, however previous sequencing of the *Drosophila* gut only identifies the presence of certain nicotinic receptor subunits and no muscarinic subunits (53, 54). Therefore, we treated the *Drosophila* midguts with nicotine, which activates nicotinic receptors without being degraded by Ace(55). Nicotine treatment increases Ca^2+^ levels in WT and to a lesser extent in CF model guts (S3D-E). This suggests that cholinergic signaling may be impaired at the receptor level in the CF model gut in addition to the increased *Ace* expression. In our snRNA-seq data set nicotinic receptors have low coverage and were therefore filtered out of the differential expression analysis (File S3). However, when the nicotinic receptors are included in the analysis, we observe a trend towards reduction in overall nicotinic receptor subunit gene expression (S3F-G). We confirmed this overall reduction in nicotinic receptor subunits by RT-qPCR (S3H). We observe a significant decrease in beta1, beta2, alpha5 and alpha7 nicotinic receptor subunits with no change in the alpha4 and alpha5 subunits and were not able to detect the alpha6 subunit mRNA via RT-qPCR (S3H). These data confirm that cholinergic signaling is reduced in the CF model by both reduced levels of nicotinic receptor subunits and by increased *Ace* expression.

To test the importance of *Ace* expression in the CF gut models sensitivity to cholinergic signaling we used the *Mex*-Gal4 (enterocyte specific driver) together with the Gal4 repressor Tubulin-Gal80^TS^ to drive expression of both *Ace^RNAi^* and *Cftr^RNAi^* in adult *Drosophila* for 2 weeks (S4). When *Ace* levels are reduced in the CF background the response to ACh is increased to WT levels (Fig 3D-E). This result indicates that reducing *Ace* expression in CF model guts is sufficient for rescuing sensitivity to cholinergic signaling even if nicotinic receptor subunit levels are also lower in the CF model. This is likely because *Ace* knockdown in the CF background reduce *Ace* levels below WT levels (S3I). Therefore, if decreased cholinergic signaling is important for CF pathophysiology reducing *Ace* expression should alter CF phenotypes in the *Drosophila* model.

### Increasing cholinergic signaling in CF model gut rescues several CF phenotypes

Recent work has demonstrated that loss of cholinergic signaling after damage leads to increased proliferation of ISCs and a failure to return to homeostasis(27). Therefore, we examined whether *Ace* expression affected the hyperplasia present in the CF model gut. Indeed, *Ace* knockdown in the CF model gut reduced proliferation in the gut indicating that decreased cholinergic signaling contributes to the increased proliferation observed in the CF intestine (Fig 4A).

**Figure 4.**
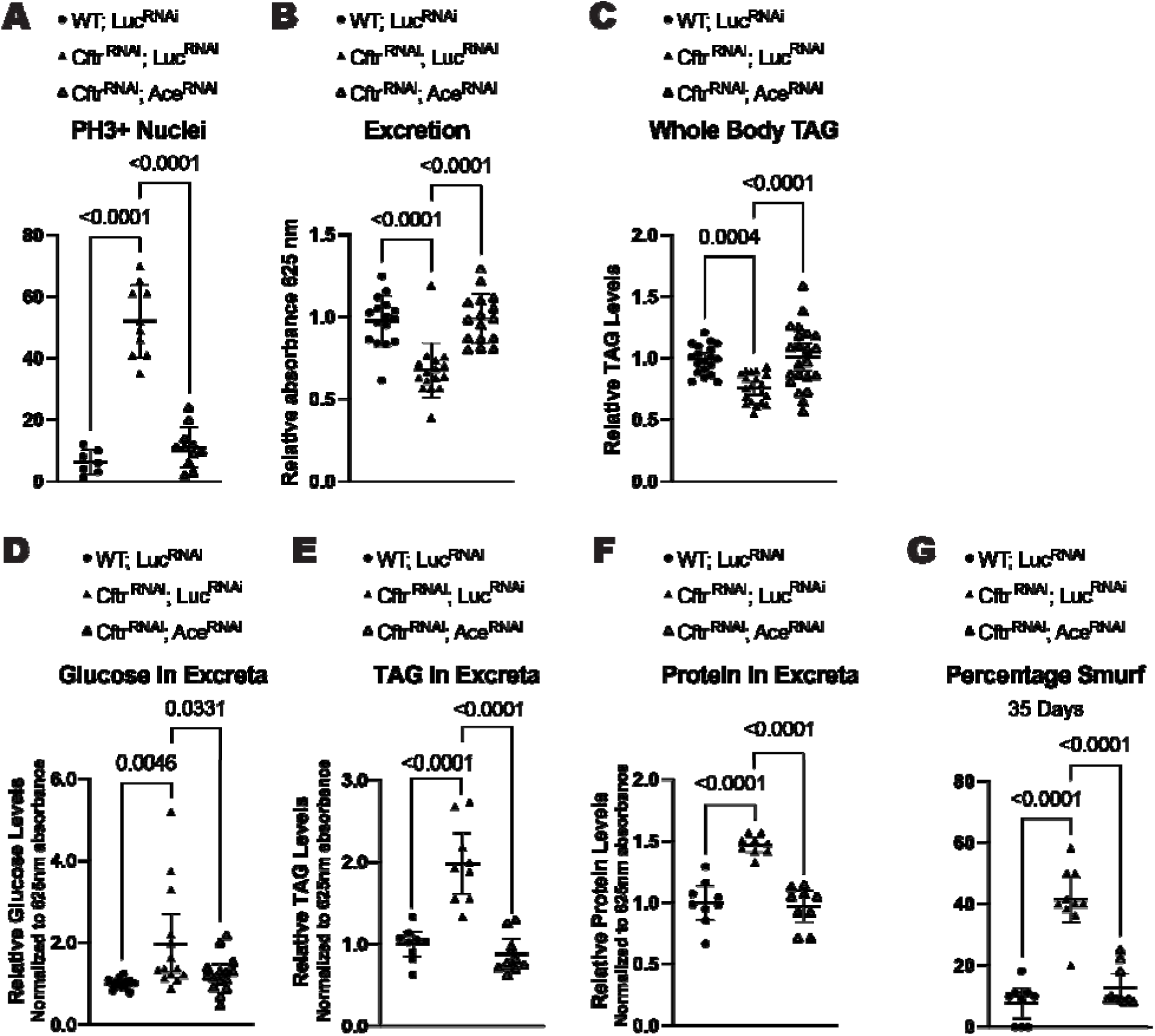
Increasing cholinergic signaling rescues CF phenotypes. (**A**) *Ace* knockdown rescues the increased proliferation seen in CF model guts. n=7 (WT; *Luc^RNAi^*), 10 (*Cftr^RNAi^*, *Luc^RNAi^*), 11 (*Cftr^RNAi^; Ace^RNAi^*). (**B**) *Ace* knockdown increases intestinal motility as measured by excretion rate in CF model guts. n=16 vials with 9-13 female flies from 3 independent experiments. (**C**) *Ace* knockdown rescues whole body TAG levels at 2 weeks of age in CF model guts. n=19 (WT; *Luc^RNAi^*), 20 (*Cftr^RNAi^*, *Luc^RNAi^*), 22 (*Cftr^RNAi^; Ace^RNAi^*) of 5 pooled females from 4 independent experiments. (**D**) *Ace* knockdown reduces the amount of glucose remaining in excreta in CF model gut flies. n=14 (WT; *Luc^RNAi^*), 14 (*Cftr^RNAi^*, *Luc^RNAi^*), 15 (*Cftr^RNAi^; Ace^RNAi^*) vials of 15-20 females from 3 independent experiments. (**E-F**) *Ace* knockdown reduces the amount of TAG (**E**) and protein (**F**) remaining in excreta in CF model gut flies. n=9 vials of 60 flies from 3 independent experiments. (**G**) CF model gut flies have increased intestinal permeability, measured by the presence of normally non-permeable blue dye in the hemolymph at 35 days of age, which is rescued by *Ace* knockdown. n=9 (WT; *Luc^RNAi^*), 10 (*Cftr^RNAi^*, *Luc^RNAi^*), 10 (*Cftr^RNAi^; Ace^RNAi^*) vials of 9-13 females from 3 independent experiments. (**A-G**) pValues were calculated using ordinary one-way ANOVA with Tukey’s multiple comparisons test in GraphPad prism. Error bars are mean with 95% CI. RNAi constructs were expressed in enterocytes using the enterocyte specific driver *Mex^TS^* (*Mex-Gal4* with the temperature sensitive gal4 repressor *tubulin gal80^TS^*) (**S4**).

We also found that increasing cholinergic signaling by reducing *Ace* levels in the CF model gut rescued the excretion rate indicating that cholinergic signaling can regulate intestinal transit in the CF model (Fig 4B). Reducing *Ace* expression in the CF model gut was also sufficient for rescuing both the whole-body TAG stores and the malabsorption of nutrients observed in the CF model gut at 2 weeks of age (Fig 4C-F).

Recently cholinergic signaling has been linked to intestinal barrier function in the *Drosophila* gut(28). As there is evidence of increased intestinal permeability in pwCF(56), we tested whether the CF model gut had decreased intestinal barrier function. To test intestinal barrier function, we performed a Smurf assay, where flies are fed a blue food dye that in healthy guts is impermeable to the intestinal barrier. When intestinal barrier function is impaired the blue dye leaks into the hemolymph and the fly turns blue (Smurf)(57). We found that CF model guts have decreased barrier function demonstrated by the increase percentage of Smurf flies at 35 days of age (Fig 4G). Importantly, intestinal barrier integrity is rescued by increasing cholinergic signally via *Ace* knockdown in the gut (Fig 4G).

Altogether these results indicate that the reduction in cholinergic signaling observed in CF model guts contributes to multiple CF pathologies.

### Fkh regulates *Ace* transcription in CF model gut

As *Ace* RNA levels are changed in the CF model gut, we performed a screen of transcription factors to identify how *Ace* expression is regulated downstream of loss of CFTR function. To be included in the screen the transcription factor had to 1) have expression in the single cell data where *Ace* is expressed (S5A) and 2) have potential binding sites as assessed by the TF2G database (Table S1)(58). We identified 18 putative transcription factors (TFs) and used RNAi to test their ability to regulate *Ace* transcription in the *Cftr* deficient background (*MyoTS*> *Cftr^RNAi^, TF^RNAi^*) by quantifying *Ace* levels (S5B). The top 2 hits with the lowest *Ace* expression in the CF background were *fork head* (*fkh*) RNAi lines (Fig 5A). In *Drosophila* Fkh has been shown to regulate ISC proliferation and expression of nutrient transporters in the intestine(59, 60). Interestingly, the mammalian ortholog FOXA1/A2 regulates expression of CFTR and other transmembrane proteins important for regulating ion homeostasis in the intestinal epithelium(35, 36). Additionally, *fkh* expression has the highest correlation with *Ace* expression in the aEC2 CF cell cluster of the transcription factors we tested (S5C). Furthermore, Fkh has 2 ChIP-Seq peaks identified at the *Ace* promoter in an embryonic ChIP-seq data set in a region identified as a cis-regulatory module (CRM)(61) for *Ace* (S5D-E). In addition to the Fkh peaks at the *Ace* promoter there are an additional 2 Fkh ChIP-seq peaks in the intronic regions of *Ace.* These intronic binding sites could also be important for regulation of *Ace* transcription especially since Fkh mammalian ortholog, FOXA1, has been shown to regulate *CFTR* expression through intronic binding sites (62–64). Together these data support but do not confirm direct regulation of *Ace* transcription by Fkh as Fkh is 1) expressed in cells that express *Ace* (S5A,C) 2) has the ability to bind to the *Ace* promoter and intronic regions (S5D-E) and 3) knockdown of *Fkh* reduces *Ace* mRNA levels (Fig 5A, S5F). Therefore, we investigated Fkh activity and function in the CF model guts.

**Figure 5.**
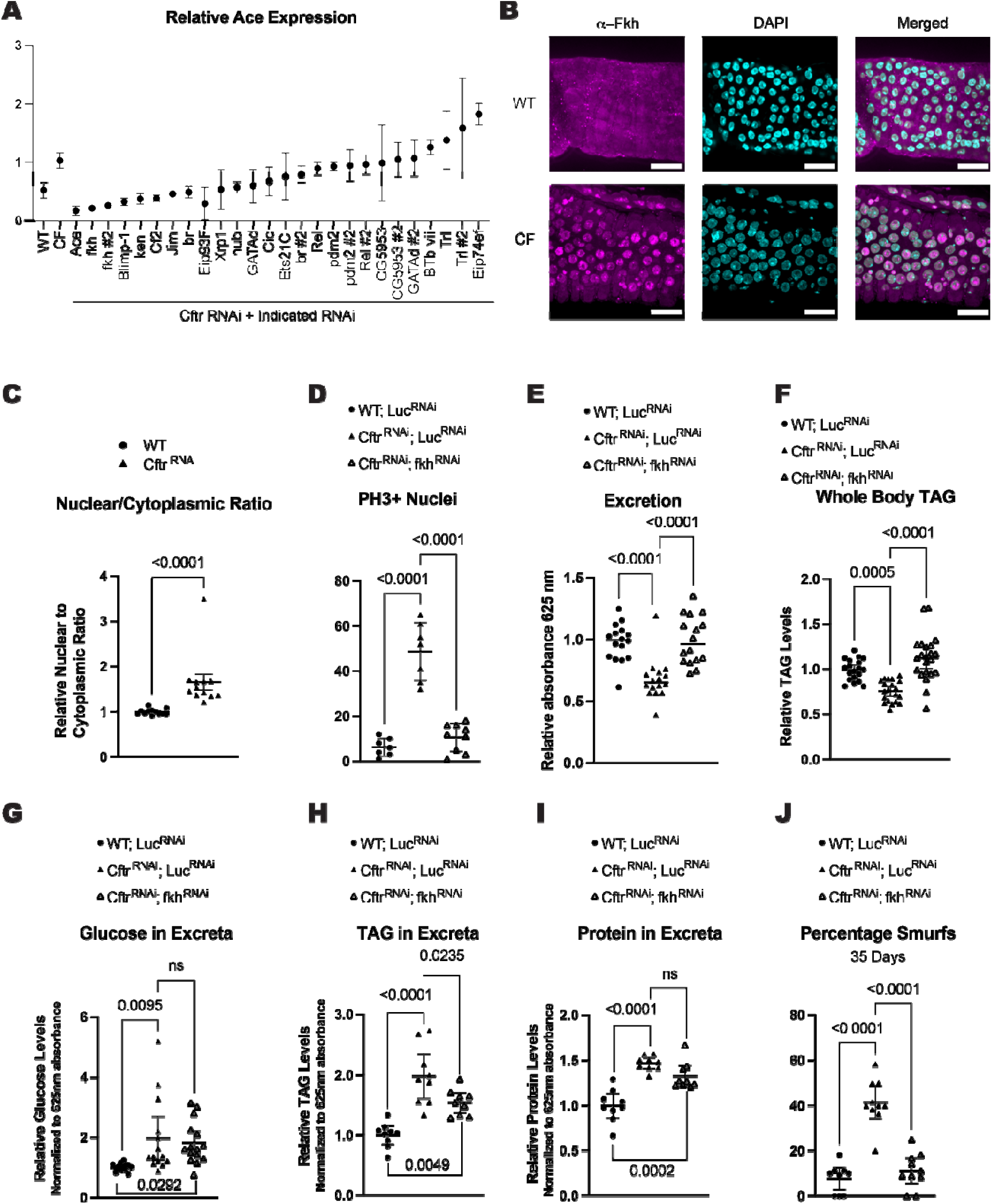
Fkh regulates *Ace* transcription and CF pathologies in the CF model gut. (**A**) Screen of transcription factors identifies Fkh as a candidate transcription factor of *Ace* in the CF model guts. n=2 biological replicates of 15 pooled guts. error bars are mean +/- SD. RNAi constructs were expressed in adult flies for 2 weeks using *Myo^TS^* (*Myo1A-Gal4, tub-Gal80^TS^)* driver (S5B). (**B**) Representative images of increased Fkh nuclear localization in CF model guts compared to WT. scale bars 25μm. (**C**) Fkh has increased nuclear localization in *Cftr* deficient guts. n=12 from 4 independent experiments. pValues were calculated using the Mann-Whitney test in GraphPad prism. Error bars are mean with 95% CI. (**D**) *fkh* knockdown rescues the hyperplasia observed in CF model guts. n= 7 (WT; *Luc^RNAi^*), 9 (*Cftr^RNAi^*, *Luc^RNAi^*), 9 (*Cftr^RNAi^*; *fkh^RNAi^*) from 2 independent experiments. (**E**) *fkh* knockdown increases intestinal motility as measured by excretion rate in CF model guts. n=16 vials with 9-13 female flies from 3 independent experiments. (**F**) *fkh* knockdown rescues whole body TAG levels at 2 weeks of age in CF model guts. n=19 (WT; *Luc^RNAi^*), 20 (*Cftr^RNAi^*, *Luc^RNAi^*), 22 (*Cftr^RNAi^*; *fkh^RNAi^*) of 5 pooled females from 4 independent experiments. (**G**) *fkh* knockdown does not reduce the amount of glucose remaining in excreta in CF model gut flies. n=14 (WT; *Luc^RNAi^*), 14 (*Cftr^RNAi^*, *Luc^RNAi^*), 15 (*Cftr^RNAi^*; *fkh^RNAi^*) vials of 15-20 females from 3 independent experiments. (**H**) *fkh* knockdown reduces TAG levels in the excreta of CF model gut flies. n=9 vials of 60 flies from 3 independent experiments. (**I**) *fkh* knockdown does not reduce the level of protein in the excreta of CF model gut flies. n=9 vials of 60 flies from 3 independent experiments. (**J**) Fkh depletion rescues intestinal barrier function in flies with *Cftr* deficient enterocytes as assessed by percentage smurf flies at 35 days of age. n=9 (WT; *Luc^RNAi^*), 10 (*Cftr^RNAi^*, *Luc^RNAi^*), 10 (*Cftr^RNAi^*; *fkh^RNAi^*) vials of 9-13 females from 3 independent experiments. (**D-J**) pValues were calculated using ordinary one-way ANOVA with Tukey’s multiple comparisons test in GraphPad prism. Error bars are mean with 95% CI. (**D-J**) Share values for WT; *Luc^RNAi^* and *Cftr^RNAi^*; *fkh^RNAi^* with Figure 4 A-E as *Ace* and *fkh* knockdown in CF background were performed at the same time and used the same controls. (**D-J**) RNAi constructs were expressed in enterocytes using the enterocyte specific driver *Mex^TS^* (*Mex-Gal4* with the temperature sensitive gal4 repressor *tubulin gal80^TS^*)(S4).

As there is minimal difference in the expression of *fkh* between WT and CF midguts in our single cell data and that Fkh activity is known to be regulated by nuclear translocation(60, 65), we examined Fkh nuclear localization. We used confocal microscopy to image the localization of Fkh in the anterior midguts, the region with highest *Ace* expression in the *Cftr* deficient guts. We found increased nuclear localization of the Fkh protein in the nuclei of CF model compared to WT guts consistent with increased Fkh transcriptional activity (Fig 5B-C, S5G-H). Fkh nuclear localization can be inhibited by phosphorylation by mTOR(60, 65). Consistent with increased Fkh nuclear localization there is decreased mTOR activity, measured by phospho-4EBP levels in anterior midguts, in the CF model gut compared to WT guts (Fig S5I-J).

If Fkh is regulating *Ace* transcription, we expect Fkh manipulation to also modulate CF pathologies. Indeed, reducing *fkh* expression lowered ISC proliferation, increased intestinal motility, increased whole body TAG stores, and increased intestinal barrier integrity in the CF model guts (Fig 5D-F,J). Interestingly, *fkh* knockdown in the *Cftr* deficient background does not fully rescue the nutrient malabsorption phenotype as shown by no change in the amount of glucose or protein remaining in the excreta and only partial reduction of the TAG levels in the excreta (Fig 5G-H). This is the one phenotype tested that does not phenocopy *Ace* knockdown, likely due to other transcriptional targets of Fkh, such as nutrient transporters, also being downregulated (60). These results suggest that increased Fkh nuclear localization downstream of CFTR loss of function regulates CF pathophysiology likely through increasing *Ace* expression.

## Discussion

We show that the *Drosophila* CF intestinal model recapitulates several clinical pathologies of CF, including poor intestinal motility, nutrient malabsorption, reduced whole body energy stores, and decreased intestinal barrier function. Additionally, we performed snRNA-seq on *Cftr* deficient and WT midguts uncovering an upregulation of *acetylcholine esterase* (*Ace*) expression in *Cftr* deficient enterocytes. This upregulation in *Ace* results in a decreased sensitivity to acetylcholine in *Cftr* deficient enterocytes. Importantly, we demonstrate that increasing cholinergic sensitivity in *Cftr* deficient enterocytes by *Ace* knockdown rescues several clinical pathologies of CF. Finally, we identify the FOXA1/A2 ortholog Fkh as a putative transcription factor for *Ace*, that has increased nuclear localization in the *Cftr* deficient midguts (Fig 6).

**Figure 6.**
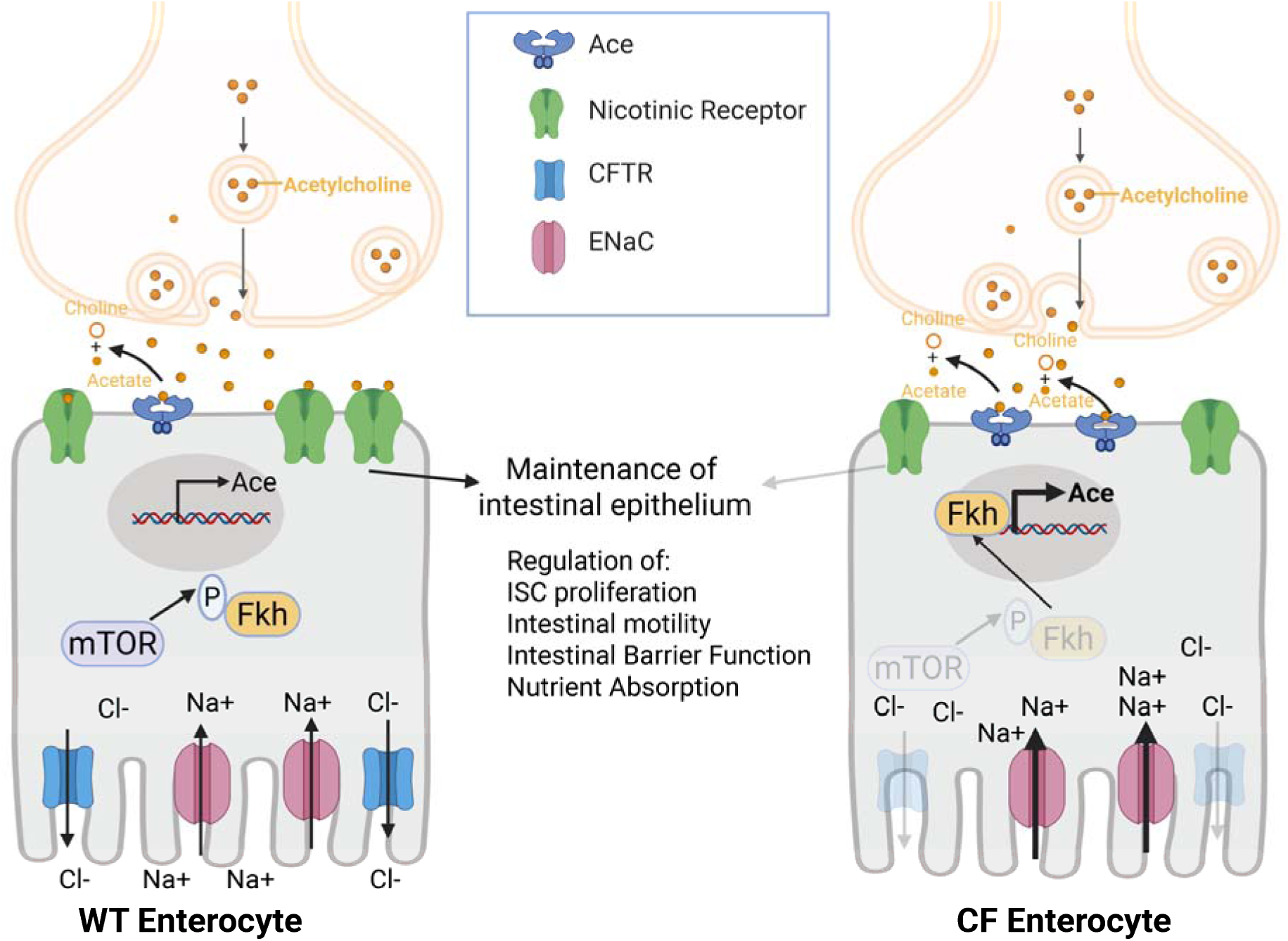
Model Figure. Loss of CFTR function in enterocytes leads to increased Fkh nuclear localization, likely from reduced mTOR activity. Increased Fkh nuclear localization increases *Ace* transcription. Increased Ace degrades acetylcholine, reducing cholinergic signaling in CF enterocytes. This reduction of cholinergic signaling impairs the maintenance of the intestinal epithelium resulting in increased ISC proliferation, decreased intestinal motility, decreased intestinal barrier function and decreased nutrient absorption. This figure was created in BioRender. Lane, L. (2025) https://BioRender.com/r4x91pu.

Previous work has shown that cholinergic signaling can increase CFTR activity(31, 32), but it has not been shown previously that CFTR can affect cholinergic signaling. Here, we demonstrate that loss of CFTR function leads to an increase in *Ace* expression and a reduction in the guts sensitivity to cholinergic signaling, indicating that there may be reciprocal regulation between cholinergic signaling and CFTR activity. This is interesting as it has been recently demonstrated in *Drosophila* that cholinergic signaling is required for a return to homeostasis after damage to the gut(27). This decrease in cholinergic signaling potential may be preventing the gut from returning to homeostasis from the damage being done by loss of CFTR. This may be because the CF gut is constantly being damaged and it may be energetically unfavorable to be constantly returning to normal homeostasis, but further studies are required to explore this hypothesis.

Additionally, we demonstrate that increasing the sensitivity to cholinergic signaling in CF model guts rescues many pathological phenotypes. Although decreasing *Ace* expression can increase cholinergic signaling in ECs it is possible that some or all of the rescue of CF pathologies with *Ace* knockdown is not cell autonomous. Ace reduces the levels of acetylcholine in the intercellular space and can affect cholinergic signaling in any nearby cell type that expresses nicotinic receptors, such as the EEs and visceral muscle. Further work will be required to definitively show which cell types are most affected by the increase in *Ace* expression in the CF model ECs. However, whether decreasing *Ace* expression in the CF model ECs is working cell autonomously or increasing cholinergic signaling in other cell types, it still improves many CF related phenotypes. Cholinergic signaling is already being investigated as a therapeutic target for diseases of intestinal inflammation such as IBD(29, 30, 66) which shares many GI symptoms with CF(67, 68). This work suggests AChE inhibition may also be therapeutically beneficial for patients with CF as well.

We also show that *Ace* transcription downstream of *Cftr* loss of function is at least in part regulated by an increase in Fkh activity. We provide evidence including Fkh ChIP-seq peaks at the *Ace* promoter and intronic region and a transcriptional response of *Ace* to Fkh depletion, that supports direct regulation of *Ace* by Fkh. However, further work such as mutational analysis would be needed to confirm direct regulation of Ace by Fkh and to fully characterize how Fkh regulates Ace transcription.

Interestingly, the human orthologs of Fkh FOXA1/FOXA2 are known to regulate the expression of CFTR as well as other ion channels important for intestinal cell function(35, 36). Therefore, in the context of ECs with decreased CFTR activity increasing FOXA1/FOXA2 activity could be a compensatory mechanism to try and increase CFTR function. The increase in Fkh activity in the CF model is not due to an increase in *fkh* transcription but instead a result of increased Fkh nuclear localization. We observe evidence of decreased mTOR activity in the CF model gut which is consistent with increased Fkh nuclear localization(60, 65). However, further studies are required to determine if the decrease in mTOR activity is cell autonomous and directly related to the loss of CFTR function. However, while Fkh activity is required for increased *Ace* transcription in the *Drosophila* CF model, it is likely not sufficient to increase *Ace* mRNA levels. Treatment of flies with rapamycin, which has been demonstrated to increase Fkh nuclear localization(60, 65), does not increase *Ace* mRNA levels after 10 or 20 days of treatment, although it is increased after 50 days, indicating that Fkh activity alone is not sufficient to increase *Ace* transcription in the midgut(69). Therefore, other transcription factors may also be important for the increased transcription of *Ace* observed in the CF model. Additional studies are needed to investigate whether any of the other transcription factors that reduced *Ace* levels in our screen are important in the context of CF. Finally, it will be important to test if this interaction between CFTR loss of function, increased Fkh activity and increased *Ace* expression is conserved in mammalian systems, as FOXA1/A2 and AChE could be therapeutically relevant targets.

We performed snRNA-Seq to compare the intestines of WT and the CF *Drosophila* model. While no unique cell clusters were identified in the CF model, we observed an increase in ISCs and newly differentiated ECs, along with a shift in EC cluster composition. The expansion of the stem cell population aligns with the increased risk of cancer in pwCF(11, 13, 39, 40). We further investigated the upregulation of *Ace* in the CF gut, prompted by recent studies highlighting the role of cholinergic signaling in maintaining intestinal epithelial homeostasis in *Drosophila*(28). In addition, we identified a substantial number of differentially expressed genes (DEGs) across all cell types, presenting numerous opportunities for future investigation. Notably, many differentially expressed secreted peptides may play a role in inter-organ communication. Together with observed changes in whole-body metabolite levels and lifespan, these findings suggest that the *Drosophila* CF model may provide valuable insights into how gut dysfunction influences systemic physiology.

Our study further establishes *Drosophila* as a useful model to study CF. Importantly, CF clinical outcomes are only partially determined by CFTR mutations and are influenced by other genetic modifiers(70–73). Many of these modifiers have been identified by GWAS studies while others are hypothesized based on interaction with CFTR or other experimental evidence(70–73). However, in most cases, these putative modifiers have not been validated and studied in detail(70–73). This work highlights the potential of *Drosophila* for investigating potential CF genetic modifiers. For example, the fly model can be used to screen which of these potential genetic modifiers alter CF pathophysiology. Understanding how CF modifiers affect physiology could lead to new drug targets that could be beneficial to all CF patients regardless of the type of CFTR mutations. Similarly, *Drosophila* would be an ideal model for a first pass to screen chemical libraries for potential CF therapeutics as a relatively cheap *in vivo* model with clinically relevant phenotypes.

## Methods

### Fly lines

The following fly stocks were used in this study: Drivers from Perrimon lab stocks: mex1-Gal4(74) (mex), mex1-Gal4 Tubulin-Gal80TS (*mex*TS), Myo31DF^NP0001^-Gal4 (myo1A-Gal4)(75), Myo1A-Gal4 Tubulin-Gal80TS(*myo1A*TS).

UAS-RNAi lines from NIG and BDSC Drosophila Stock Centers: UAS-Cftr^RNAi^- NIG Stocks: 5789R-1 & 5789R-4, UAS-Ace^RNAi^- BDSC 25958, UAS-Luciferase^RNAi^-BDSC 31603 UAS-fkh^RNAi^- BDSC 27072, UAS-fkh^RNAi^ #2- BDSC 58059, UAS-Blimp-1^RNAi^- BDSC57479, UAS-Ken^RNAi^- BDSC-34739, UAS-CF2^RNAi^- BDSC 57256, UAS-Jim^RNAi^- BDSC 42662, UAS-br^RNAi^-BDSC 33641, UAS-Eip93F^RNAi^ BDSC 57868, UAS-Xrp1^RNAi^- BDSC 51054, UAS-nub^RNAi^-BDSC 28338, UAS-GATAd^RNAi^- BDSC 34640, UAS-Cic^RNAi^- BDSC 25995, UAS-Ets21c^RNAi^-BDSC 39069, UAS-br^RNAi^ #2- BDSC 27172, UAS- rel^RNAi^- BDSC 28943, UAS-pdm2^RNAi^-BDSC 29453, UAS-pdm2^RNAi^ #2- BDSC 50665, UAS-rel^RNAi^ #2- BDSC 35661, UAS-CG5953^RNAi^- BDSC 57543, UAS-CG5953RNAi #2- BDSC 57287, UAS-GATAd^RNAi^-BDSC 33747, UAS-BTb-viiRNAi- BDSC 28912, UAS-Trl^RNAi^- BDSC 40940, UAS-TrlRNAi #2-BDSC 41582, UAS-Eip74ef-BDSC 29353.

### Excretion assay

Flies were fed overnight on lab food with 2 g/100 ml FD&C Blue Dye (12-15 flies per vial for female and 15-20 flies per vial for male). After overnight feed they were moved to 5 ml culture tube without food for 2 hrs (female) or 1 hr 45 minutes (males), time points where the flies have not completely cleared the food from their midguts. Excreta was collected in 400 ul of .05% PBST and 100 ul was used in triplicate to measure blue absorbance at 625 nm using the SpectraMax Paradigm Multi-mode microplate reader (Molecular Devices). A standard curve was made with serial dilutions of the FD&C Blue Dye in .05% PBS and values were normalized to WT for each experiment.

### Whole body metabolite measurements

For each replicate 5 flies (females) or 8 flies (males) were collected in eppendorph tubes containing 1.0 mm Zirconium Oxide Beads (Next Advance Lab Products—ZROB10) and frozen on dry ice and stored at −80C. Samples were moved to ice and 500 μL of ice-cold PBS with 0.05% Triton-X-100(Sigma) was added to the tubes. Homogenization was carried out using a TissueLyser II (QIAGEN) for 2-3 cycles of 30 seconds each, at an oscillation frequency of 30 Hz/s. Tubes were centrifuged for 1 minute at 3500 g to remove debris, and the homogenate was immediately used for glucose, protein, and TAG quantification. Quantifications were performed using a SpectraMax Paradigm Multi-mode microplate reader (Molecular Devices). For protein measurements, 5 μL of the homogenate was mixed with 200 μL of the BCA Protein Assay Kit in triplicate (Pierce BCA Protein Assay Kit), followed by incubation for 30 minutes at 37°C with gentle shaking in 96-well Microplates (Greiner Bio-One) and measuring absorbance at 562nm. Glucose quantification involved mixing 10 μL of homogenate with 100 μL of Infinity Glucose Hexokinase Reagent in triplicate (Thermo Fisher Scientific—TR15421), incubating for 30 minutes at 37°C in 96-well Microplates UV-Star (Greiner Bio-One – 655801) with gentle shaking and measuring 340nm. For triglycerides, 5 μL of the homogenate was mixed with 150 μL of Triglycerides Reagent in triplicate (Thermo Fisher Scientific—TR22421) and incubated for 10 minutes at 37°C in 96-well Microplates (Greiner Bio-One) with gentle shaking and absorbance was measured at 520 nm. Values were calculated from standard curves using seven serial dilutions (1:1) of BSA (Pierce BCA Protein Assay Kit), glycerol standard solution (Sigma), or glucose standard solution (Sigma). Glucose and TAG levels were normalized to BCA protein levels and then normalized to WT levels for each independent cross.

### Glucose in excreta

Flies were fed overnight on lab food supplemented with 1 g/ L FD&C Blue dye with 15-20 flies per vial. Flies were transferred to 5 ml culture tubes with 200 ul of lab food supplemented with 1 g/ L FD&C Blue Dye for 6 hr. Excreta was collected in 200 ul of .05% PBST and transferred to a 96 well microplate (Greiner Bio-One – 655801) and absorbance was measured at 625 nm using the SpectraMax Paradigm Multi-mode microplate reader (Molecular Devices). A standard curve was made with serial dilutions of the FD&C Blue Dye in .05% PBS. Glucose quantification was performed as in whole body metabolite measurements and glucose levels were normalized to total excreta amount (625 nm) absorbance before being normalized to the WT condition in each experiment.

### TAG and Protein in Excreta

Flies were fed overnight on lab food supplemented with 1 g/ L FD&C Blue dye with 60 flies per bottle. Flies were transferred to 5 ml culture tubes with 200 ul of lab food supplemented with 1 g/ L FD&C Blue Dye for 4 hr. Excreta was collected in 200 ul of .05% PBST and transferred to a 96 well microplate (Greiner Bio-One – 655801) and absorbance was measured at 625 nm using the SpectraMax Paradigm Multi-mode microplate reader (Molecular Devices). A standard curve was made with serial dilutions of the FD&C Blue Dye in .05% PBS. TAG quantification was performed as in whole body metabolite measurements section. Protein levels were measured via BCA assay as described in whole body metabolite measurements section. TAG and protein levels were normalized to total excreta amount (625 nm) absorbance before being normalized to the WT condition in each experiment.

### Feeding Assay

#### 24hr. Feeding Assay

The amount of food consumed over a 24hr period was measured using the DIETS assay (76). 25 flies were place in a vial with agar plug (as water source) and a small amount of food (100ul) was placed in an eppendorph tube cap. The cap with food was affixed with tape to the side of the fly vial. The flies were acclimatized to the system for 24hrs before being transferred to a new vial with agar plug with a weighed eppendorph tube cap with 100 ul of food. After 24 hrs of consumption the food was weighed again. Empty vials with food caps were used as evaporation controls. Food intake per fly was calculated as (change in weight-evaporation controls change in weight)/ number of flies.

### 30 minute blue dye feeding assay

Flies were starved overnight before being put on standard lab food containing 1g/100ml FD and C blue dye for 30 minutes. 5 flies with heads removed were homogenized in 600ul of .05% PBST and tubes were were centrifuged for 1 minute at 3500 g to remove debris. 100 ul was used in triplicate per replicate to measure blue absorbance at 625 nm using the SpectraMax Paradigm Multi-mode microplate reader (Molecular Devices). A standard curve was made with serial dilutions of the FD&C Blue Dye in .05% PBS and values were normalized to WT for each experiment.

### Lifespan assay

15 (females) or 20 (males) 2-4 day old adult flies were placed in vials on standard laboratory food. Flies were flipped every other day and vials were scored for percent survival.

### Smurf assay

Smurf assay was performed as previously described(57). Briefly 10-15 female flies were raised on standard fly food and flipped every 2 days for 33 days before being put on standard fly food supplemented with 2.5 g per 100 ml of FD & C blue dye for 2 days. At 35 days each vial was scored for percentage smurf flies under a dissection microscope.

### RNA isolation, RT and quantitative PCR

For RNA isolation, 10-15 adult guts were dissected, and tissues were homogenized in TRIzol reagent (Ambion). The supernatant was collected and processed using Direct-zol RNA MicroPrep columns (Zymo Research) following the manufacturer’s protocol. Reverse transcription was performed using the iScript cDNA Synthesis Kit (Bio-Rad). Quantitative real-time PCR (qRT-PCR) was carried out on a CFX96 Real-Time System (Bio-Rad) using iQ SYBR Green Supermix (Bio-Rad). qRT-PCR reaction volume used was 10 µl (2 µl 5 µM Primer pair mix+ 5 µ 2x SYBR Green+ 3 µl cDNA). Relative mRNA levels were determined using the ΔΔCt method, with mRNA levels normalized to Rpl32.

qPCR primers:

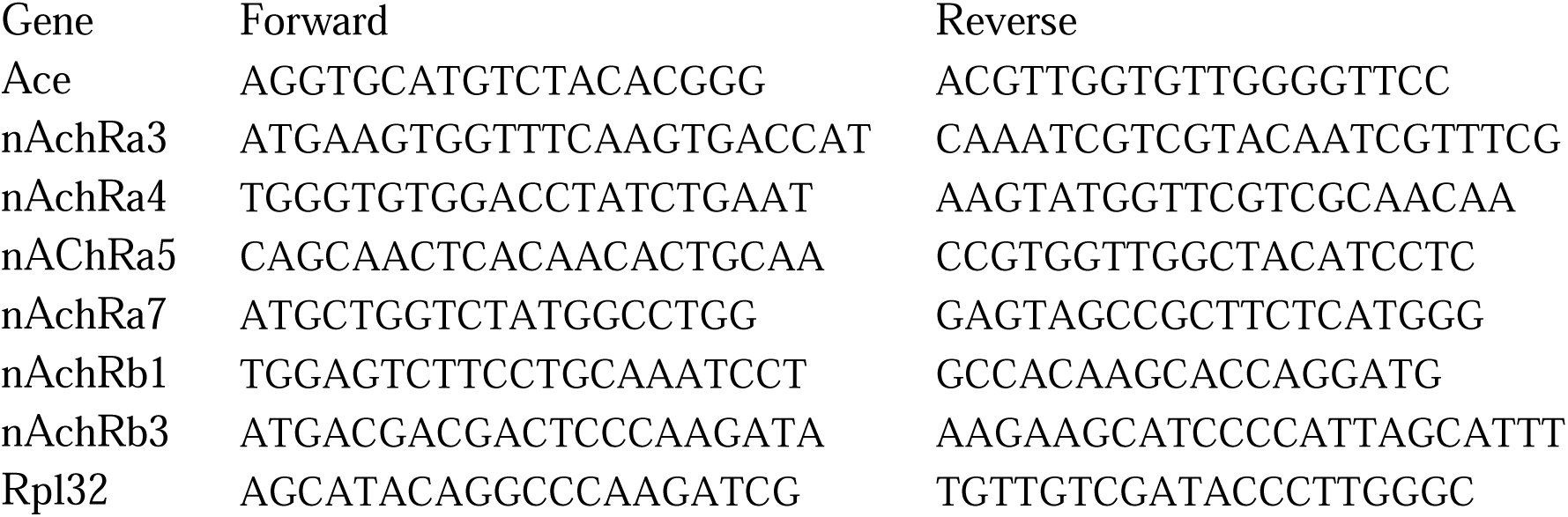

### Single-nuclei RNA-seq

#### Single-nucleus suspension and FACS

Single-nucleus suspension was conducted as previously described(47). Briefly, 70 guts per condition were dissected in cold Schneider’s medium, flash-frozen and stored at −80°C. Prior to FACs sorting, samples were spun down and Schneider’s medium was exchanged with homogenization buffer [250mM Sucrose, 10mM Tris pH8, 25mM KCl, 5mM MgCl, 0.1% Triton-X, 0.5% RNasin Plus (Promega, N2615), 50x protease inhibitor (Promega G6521), 0.1mM DTT]. Using 1ml dounce (Wheaton 357538), nuclei were released by 20 loose pestle strokes and 40 tight pestle strokes while keeping samples on ice and avoiding foam. Next, nuclei were filtered through 5 ml cell strainer (40 μm), and using 40 μm Flowmi (BelArt, H13680-0040). Nuclei were centrifuged, resuspended in PBS/0.5%BSA with 0.5% RNase inhibitor, filtered again with 40 μm Flowmi and stained with DRAQ7 Dye (Invitrogen, D15106). Single nuclei were sorted with Sony SH800Z Cell Sorter at PCMM Flow Cytometry Facility at Harvard Medical School and 100k nuclei per sample were collected in PBS/BSA buffer.

#### 10x genomics and sequencing

Single nuclei RNA-seq libraries were prepared using the Chromium Next GEM Single Cell 3’ Library and Gel Bead Kit v3.1 according to the 10xGenomics protocol. Approximately 16,500 nuclei were loaded on Chip G with an initial concentration of 700 cells/µl based on the ‘Cell Suspension Volume Calculator Table’. Sequencing was conducted with Illumina NovaSeq 6000 at Harvard Medical School Biopolymers Facility.

#### 10x data processing

Raw sequencing data were aligned to the *Drosophila melanogaster* reference genome (BDGP6.32, Ensembl release 104) using 10x Genomics Cell Ranger v7.1.0. Low quality nuclei with less than 500 Unique Molecular Identifiers (UMI) were filtered out. The default clustering and uniform manifold approximation and projection (UMAP) generated by 10X Loupe Browser was used to cluster cells. Cell types were manually annotated using the top marker genes from each cluster. The filtered count matrix and metadata were loaded into R (v4.4.0) and further processing was performed using the default Seurat workflow (v5.2.1). Differential gene expression analysis was performed using a Mann Whitney-U test with a cut-off of p-value < 0.05 and abs(Log2FC) > .5 and percent expression > 1%.

Normalized expression violin plots and heatmaps were generated using Seurat, ggplot2 (v3.5.2), ComplexHeatmap (2.24.0) and pheatmap (v1.0.12). hdWGCNA (v0.4.05) was used to create metacells for which Pearson correlation tests were performed for candidate transcription factors and Ace expression.

### Cell type comparison between *Drosophila* and mammalian intestinal epithelium

We used DIOPT (release 7, score>3) (77) to map mouse genes to *Drosophila* orthologs. The *Drosophila* orthologs of markers identified in various cell types from mammalian datasets were grouped respectively as cell type specific gene sets. Next, we compared the cell type specific gene sets from the mammalian study (48) and the cell type markers from *Drosophila* (*46*). P-value of enrichment was calculated based on the hypergeometric distribution and the similarity of Drosophila markers with mammalian markers is reflected by the negative log10 of the P values.

### GCaMP7c live imaging

GCaMP7c live imaging was performed as previously described(27). Guts were dissected in fresh HL3 buffer (1.5mM Ca2+, 20mM MgCl2, 5mM KCl, 70mM NaCl,10mM NaHCO3, 5mM HEPES, 115mM Sucrose, 5mM Trehalose) and placed in eight-well clear bottom cell culture chamber slides with HL3. Guts were stabilized with a Nylon mesh (Warner instruments, 64-0198) and paper clips cut in small identical pieces. ACh and Nicotine sensitivity assay: ACh and Nicotine sensitivity assay was done using LSM780 and LSM980 microscopes with 40x water objective lens. Each frame (∼2.5 sec/frame) is the maximum projection of 5-6 z-stacks (2.96µm/z) and was acquired with 488nm excitation for GFP. 5mM Acetylcholine (Acetylcholine Chloride, Sigma A2661) or 1.33 mM Nicotine (Sigma, N3876) were added at the 10th frame (∼25 sec) For Figure 3B-C an old aliquot of Acetylcholine was used at 10mM after testing response in WT midguts to a range of concentrations, when new aliquot was purchased 5mM concentration was used after testing a range of concentrations in WT guts as giving a similar response to previous experiment. All images were taken from similar areas in the Anterior midgut between R1-R2. Fiji was used for assembly and calculation of fluorescence per frame. DF/F0 = Ffr-F0/ F0. Ffr is the fluorescence per frame and F0 (baseline fluorescence) is the average fluorescence intensity of the first 9 frames (fr1-fr9).

### PH3+ counts

Guts were dissected and fixed in 4% PFA for 20 minutes. They were washed 3x in .1% PBST and incubated for 1 hr at RT in 5% Normal Goat Serum in .1%PBST. Guts were incubated for overnight with rabbit anti-pH3 (Millipore #06-570; 1:3000). Guts were then washed for 30 minutes 3x in .1% PBST then incubated for 2hrs at RT with secondary antibody, Alexa Fluor 555-conjugated goat anti-rabbit IgG (1:200) and DAPI (1:3000). The guts were then washed 3X in .1%PBST for 30 minutes and mounted in VECTASHIELD antifade mounting media (Vector Laboratories). Slide genotype was blinded and PH3+ nuclei were counted with an epifluorescence microscope.

### Fkh nuclear localization

Guts were dissected and fixed in 4% PFA for 20 minutes. They were washed 3x in .1% PBST and incubated for 1 hr at RT in 5% Normal Goat Serum in .1%PBST. Guts were incubated for 2 days at 4C with Rab-anti-FKH (60, 78–80) (1:100)(gift from Dr. Chandrasekaran, St. Mary’s College of California) in 5% Normal Goat Serum in .1%PBST. Guts were then washed for 30 minutes 3x in .1% PBST then incubated overnight in secondary antibody, Alexa Fluor 555-or 647 conjugated goat anti-rabbit IgG (1:200) and DAPI (1:3000). The guts were then washed 3X in .1%PBST for 30 minutes and mounted in antifade mounting media. Fluorescence was imaged using W1 Yokogawa spinning disk, Nikon inverted Ti2 confocal microscope. Percent nuclear localization was calculated on maximum intensity projections in Fiji.

### Phospho-4EBP1 Staining

Guts were dissected and fixed in 4% PFA for 30 minutes. They were washed 3x in .1% PBST and incubated for 1 hr at RT in 5% Normal Goat Serum in .1%PBST. Guts were incubated for 2 days at 4C with Rab-anti-p4EBP1 (1:800) (CST 2855S) in 5% Normal Goat Serum in .1%PBST. Guts were then washed for 30 minutes 3x in .1% PBST then incubated overnight in secondary antibody, Alexa Fluor 555-or 647 conjugated goat anti-rabbit IgG (1:200) and DAPI (1:3000). The guts were then washed 3X in .1%PBST for 30 minutes and mounted in antifade mounting media. Fluorescence was imaged using W1 Yokogawa spinning disk, Nikon inverted Ti2 confocal microscope. Average staining intensity was calculated on maximum intensity projections in Fiji.

### Statistics and Reproducibility

Prism (https://www.graphpad.com/scientific-software/prism/) or ggplot2 was used to create graphs as well as to perform statistical analysis. Statistical tests are indicated in figure legends. No statistical analysis was conducted to determine sample sizes. Randomization was not performed. Blinding experiments were conducted during mitotic division counts. No data were excluded. Confocal images shown in figures are representative of two to 4 independent experiments with similar result.

## Data Sharing/ Resource Availability

The scRNA-seq portal for data mining is available at https://www.flyrnai.org/scRNA/. Raw RNA-seq reads and processed data files have been deposited in the NCBI Gene Expression Omnibus (GEO) database under accession code: GSE300597.

## Author Contributions

Conceptualization: Elizabeth A. Lane, Norbert Perrimon

Data Curation: Yanhui Hu, Weihang Chen, Mujeeb Qadiri

Formal Analysis: Elizabeth A. Lane, Afroditi Petsakou, Mujeeb Qadiri

Funding Acquisition: Elizabeth A. Lane, Norbert

Perrimon Investigation: Elizabeth A. Lane, Afroditi Petsakou, Ying Liu

Methodology: Elizabeth A. Lane, Afroditi Petsakou, Ying Liu

Project Administration: Elizabeth A. Lane, Norbert Perrimon

Software: Weihang Chen, Mujeeb Qadiri, and Yanhui Hu

Supervision: Norbert Perrimon

Visualization: Elizabeth A. Lane, Ying Liu, Mujeeb Qadiri, and Yanhui Hu

Writing – Original Draft Preparation: Elizabeth A. Lane

Writing – Review & Editing: Norbert Perrimon, Afroditi Petsakou, Ying Liu

This article is subject to HHMI’s Open Access to Publications policy. HHMI lab heads have previously granted a nonexclusive CC BY 4.0 license to the public and a sublicensable license to HHMI in their research articles. Pursuant to those licenses, the author-accepted manuscript of this article can be made freely available under a CC BY 4.0 license immediately upon publication.

## Acknowledgments

Confocal imaging was conducted at MicRoN Facility at Harvard Medical School, and we thank Paula Montero Llopis for advice. We also thank Biopolymers Facility and Computing facilities and PCMM Flow Cytometry Facility at Harvard Medical School. We thank Sudhir Gopal Tattikota for his expertise on 10x snRNA-seq. We thank Muhammad Ahmad for advice on image analysis. We thank DRSC/TRiP, NiG and the Bloomington Stock Center for fly lines. We thank Dr. Chandrasekaran for providing the anti-Fkh antibody.

## Declaration of generative AI and AI-assisted technologies in the writing process

During the preparation of this work the author(s) used ChatGTP in order to polish certain sections of text. After using this tool/service, the author(s) reviewed and edited the content as needed and take(s) full responsibility for the content of the publication.

## Supplementary Materials

S1 Appendix: Supplementary Figures S1-S5 and Supplementary Figure Legends

Table S1: Supplementary Table 1- TF2TG

File S1: Differentially Expressed Genes in sn-RNA-seq

File S2: Differentially Expressed Secreted Peptide Genes in sn-RNA-seq

File S3: Average Expression Nicotinic Receptors in sn-RNA-seq

Data S1: Numerical Data Underlying Main Figure Graphs

Data S2: Numerical Data Underlying Supplementary Figure Graphs

