## Supplementary material for "Cholinergic Signaling Modulates Intestinal Pathophysiology in a *Drosophila* Model of Cystic Fibrosis": S1 Appendix

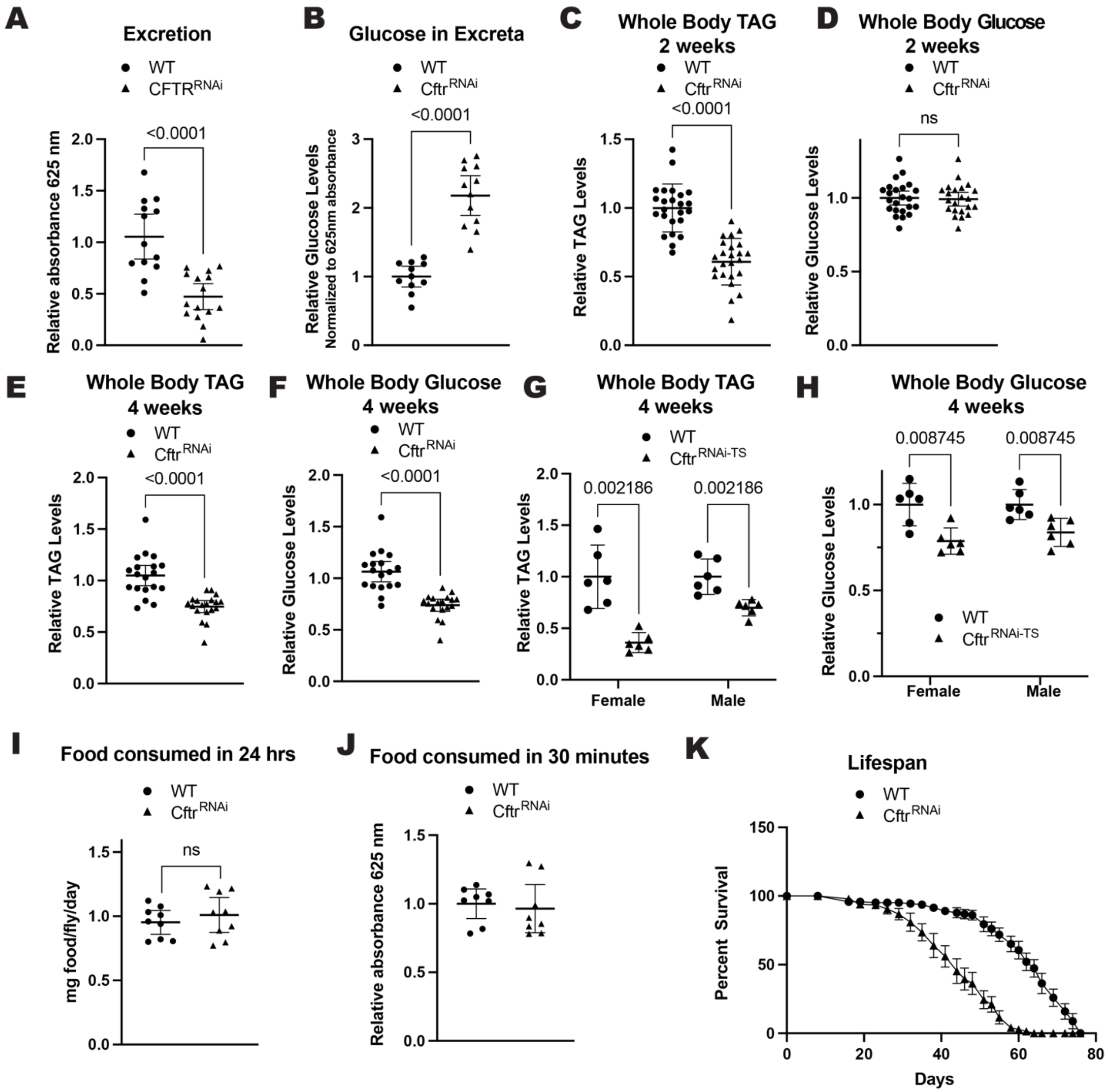


**S1. Related to Figure 1**

(**A**) CF model guts have decreased excretion rate compared to WT guts as measured by the amount of excreta collected over a 1.75 hr time period. n= 13(WT), 15 ($Cftr^{RNAi}$) vials of 10-15 males from 3 independent crosses. (**B**) CF model guts have increased glucose in excreta compared to WT flies. N= 11 (WT), 12 ($Cftr^{RNAi}$) vials of 15-20 males from 2 independent crosses. (**C-F**) CF model guts have reduced whole body energy stores to WT flies. (**C**) CF model guts have reduced TAG levels at 2 weeks of age compared to WT flies. n= 24 of 8 pooled males from 4 independent crosses. (**D**) Male CF model gut flies have no significant difference in whole body glucose levels at 2 weeks of age compared to WT flies. n= 23 (WT), 22 ($Cftr^{RNAi}$) 8 pooled males from 4 independent crosses. (**E**) CF model gut flies have reduced whole body TAG levels at 4 weeks of age compared to WT flies. n= 19 of 8 pooled males from 4 independent crosses. (**F**) CF model gut flies have reduced whole body glucose at 4 weeks of age compared to WT flies. n= 24 (WT), 23 (*Cftr*$^{RNAi}$) of 8 pooled males from 4 independent crosses. (**G-H**) Decreased whole body metabolites are not due to a developmental defect in CF model guts. (**G**) Whole body TAG levels are reduced in CF model guts compared to WT when *Cftr* knockdown is induced by temperature shift in 1-2 day old adults. n= 6. **(H)** Whole body glucose levels are reduced in CF model guts compared to WT when *Cftr* knockdown is induced by temperature shift in 1-2 day old adults. n= 6 of 5 (female) or 8 (male) pooled flies. (**I**) CF model gut flies eat a similar amount of food over a 24 hr period as WT flies. n=9 vials with 25 flies from 2 independent experiments. (**J**) CF model gut flies eat a similar amount of food in the 30 minutes after starvation as WT flies. n=8 of 5 pooled females. (**A-J**) pValues were calculated using the Mann-Whitney test in Graphpad prism. Error bars are mean with 95% CI. (**K**) CF model gut flies have reduced lifespan compared to WT flies. n= 11 vials with 10-15 males from 2 independent crosses. (**A-F, I-K**) WT is *Myo1A* > + and *Cftr*$^{RNAi}$ is *Myo1A* > *Cftr*$^{RNAi}$. (**G-H**) WT is *Myo*$^{TS}$ > + and *Cftr*$^{RNAi\text{-}TS}$ is *Myo*$^{TS}$ > *Cftr*$^{RNAi}$.

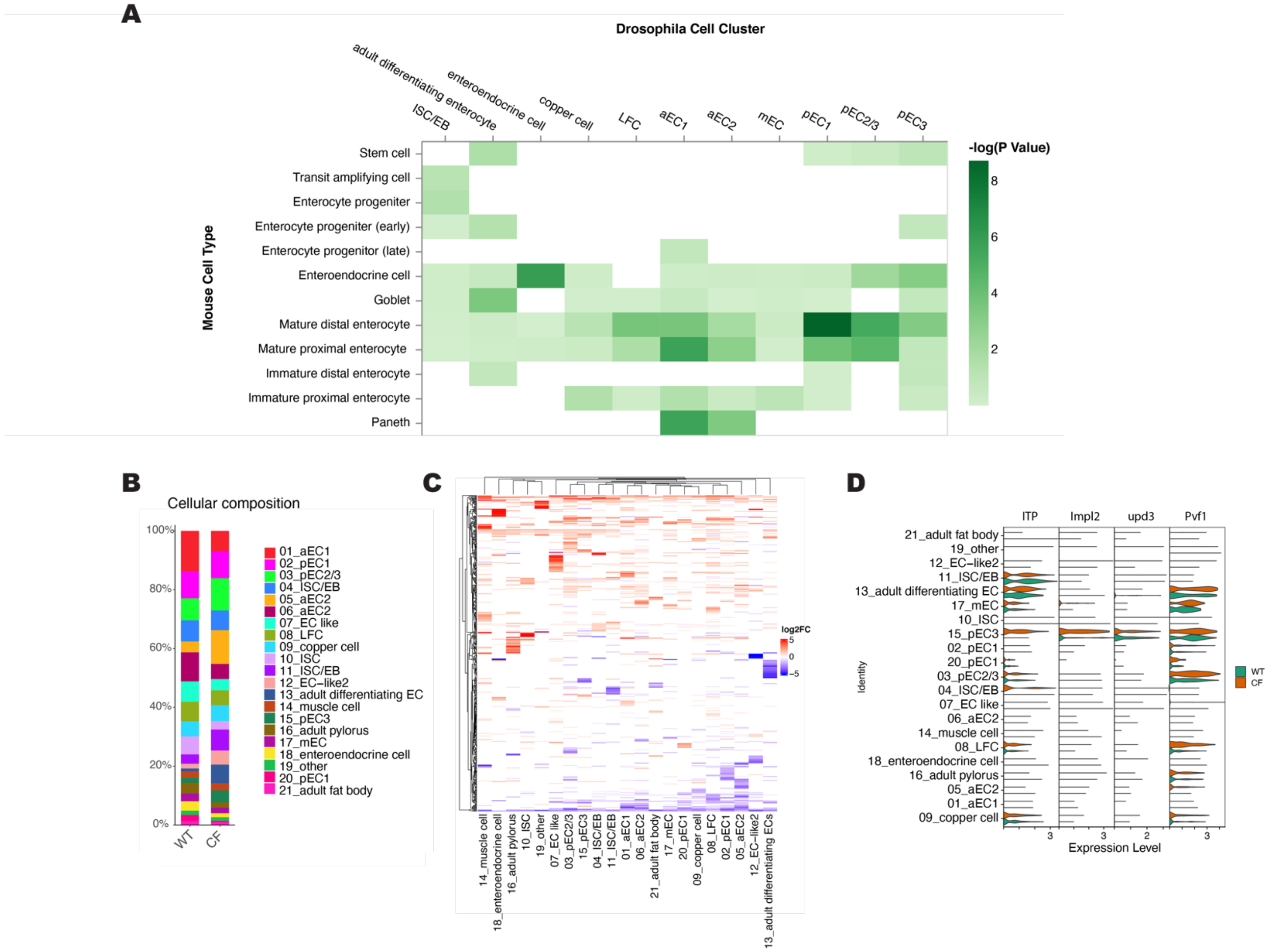


**S2. Related to Figure 2**

**(A)** Comparison of fly gut cell types and mammalian intestine cell types using marker genes identified. **(B)** Differences in cell type composition between WT and CF guts illustrated as a stacked bar chart depicting percentage of cell belonging to each cell cluster in snRNA-seq data for WT and CF model guts. **(C-D)** Many secreted peptides are differentially expressed between WT and CF guts across cell clusters. **(C)** Heat map of the differentially expressed secreted proteins. Color reflects the log2fold change of expression in CF model guts comparing to control in each cell type. Gene names of differentially expressed genes can be found in File S2. **(D)** Violin plots of expression of secreted peptides important for Yki$^{act}$ gut tumor physiology (*Itp, Impl2, upd3,* and *pvf1*) in snRNA-seq data.

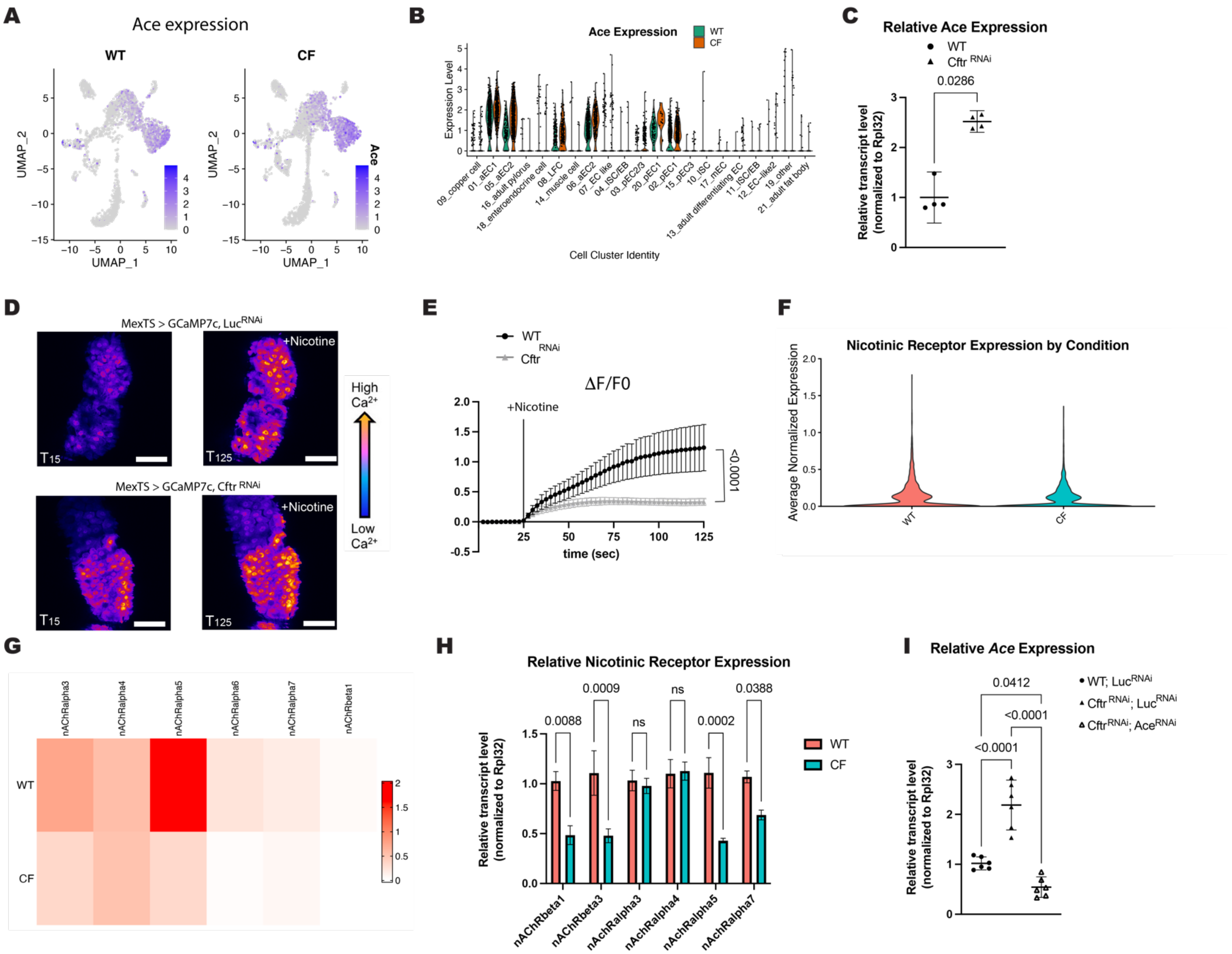


**S3. Related to Figure 3**

**(A-B)** *Ace* expression is upregulated *Cftr* deficient midguts compared to WT midguts in snRNA-seq data. **(A)** *Ace* expression in each cell in WT and *Cftr* deficient guts plotted onto UMAP **(B)** Violin plots of *Ace* expression in each cell cluster identified in snRNA-Seq data set. **(C)** *Ace* expression is increased in *Cftr* deficient guts compared to WT guts via RT-qPCR analysis of whole guts. n=4 of 10-15 pooled guts from 1 independent experiment. **(D)** Representative images of GCaMP7c fluorescence, in WT (*MexTS > GCaMP7c, LucRNAi*) and *Cftr* deficient guts (*MexTS > GCaMP7c, $Cftr^{RNAi}$*) before (T15s) or after addition of Nicotine (T125). Scale bars are 50 µm. **(E)** Graph of average relative fluorescent intensity, ΔF/F0, per frame (2.5s per frame) and genotype. n= 8 (WT) and 7 (*$Cftr^{RNAi}$*) from 3 independent experiments. Error bars are mean +/- SEM and pValue was calculated using the Mann-Whitney test in Graphpad prism. (**F**) Average normalized expression of all nicotinic receptors in WT and CF single nuclei RNA-seq data set. (**G**) Heat map of average expression of nicotinic receptor subunits in WT and CF midguts from snRNA-seq data set. (**H**) Relative expression of indicated nicotinic receptor subunit in WT and

CF model guts as assessed by RT-qPCR. n= 6-7 replicates of 10 pooled guts from 2 independent experiments. pValues were calculated using ordinary one-way ANOVA with Tukey's multiple comparisons test in GraphPad prism. Error bars are mean +/- SD (**I**) Relative *Ace* expression in guts of indicated genotype. n=6 replicates of 10 pooled guts from 2 independent experiments. pValues were calculated using ordinary one-way ANOVA with Tukey's multiple comparisons test in GraphPad prism. Error bars are mean with 95% CI.

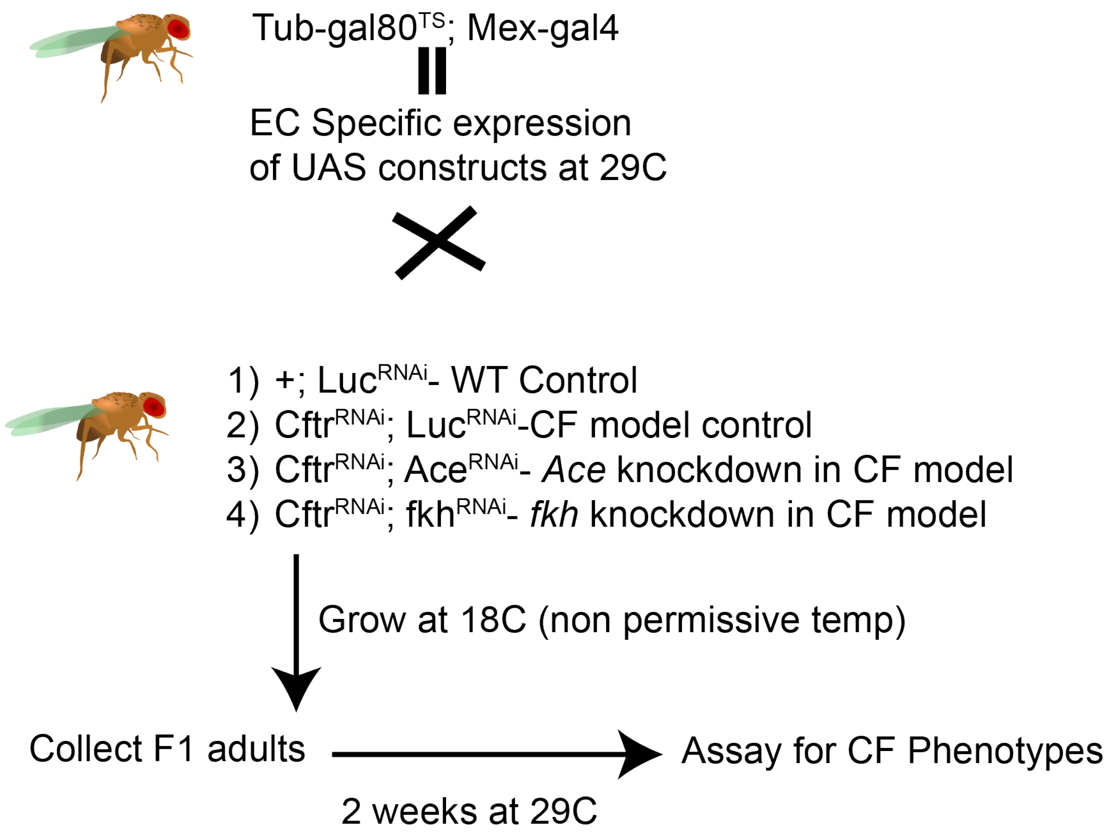


**S4. Related to Figure 3-5**

Experimental set up for figures 3D-E, 4, and 5D-H. *MexTS* (*Tub-gal80$^{TS}$, Mex-gal4*) virgin females were crossed to males with the indicated UAS-RNAi constructs and raised at 18C, nonpermissive temperatures. 1-3 day old adults were moved to 29C for 2 weeks (35 days for smurf assay) and were than used in indicated assays. *Drosophila* image from bioicons licensed under CC0.

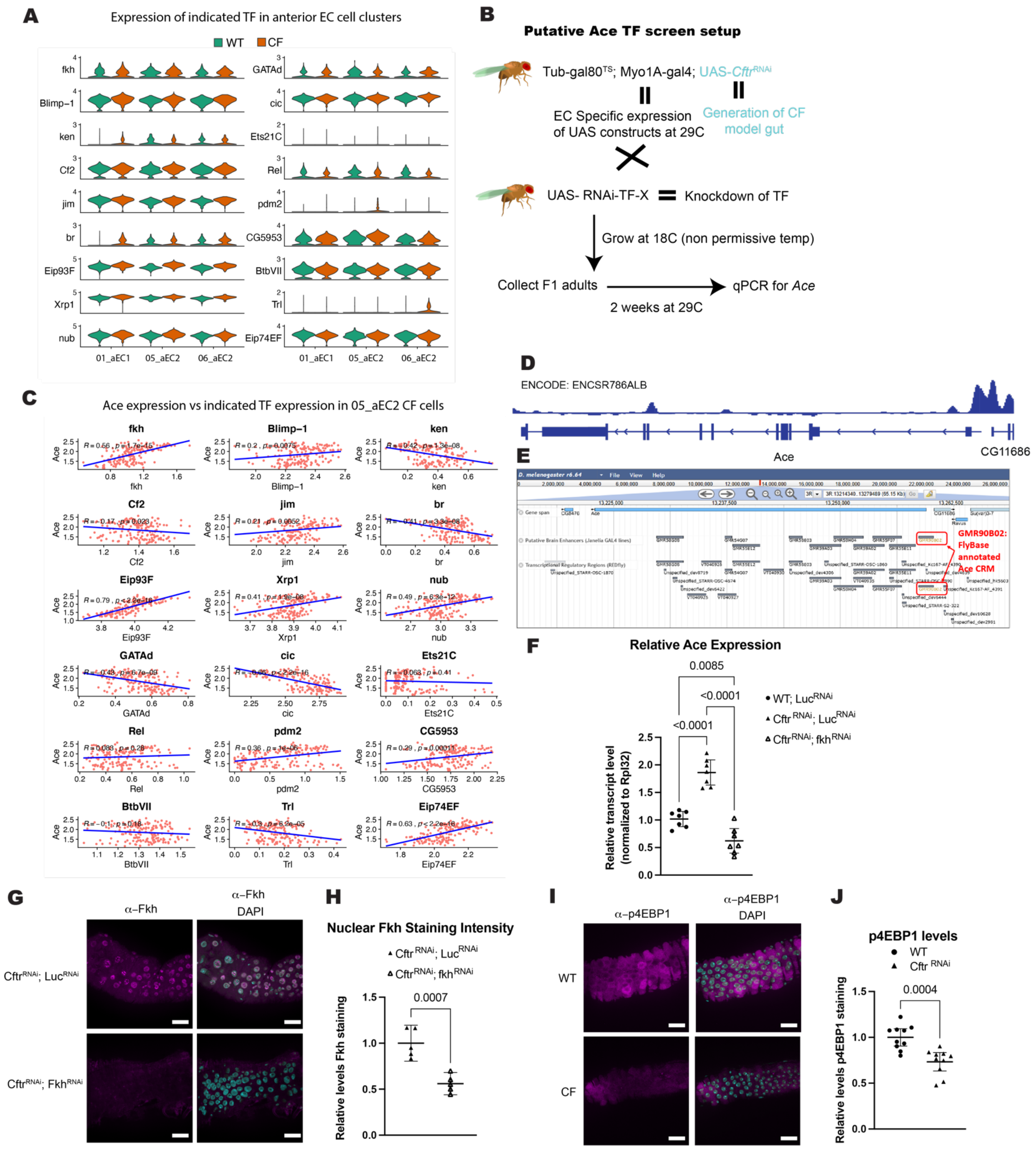


74

75 **S5. Related to Figure 5**

**(A)** Violin plots of expression of candidate *Ace* transcription factors in anterior EC cell clusters from snRNA-seq data. **(B)** Diagram of experimental set up for *Ace* transcription factor screen. *MyoTS* (*Tub-gal80TS, Myo1A-gal4*); *UAS-Cftr*$^{RNAi}$ females were crossed to male flies with UAS-RNAi against candidate transcription factor and raised at 18C, nonpermissive temperatures. 1-3 day old adults were moved to 29C for 2 weeks and guts were dissected to assay for *Ace* expression levels. **(C)** Scatter plots showing correlation of *Ace* and candidate transcription factor expression levels in meta-cells from the 05-aEC2 CF cell cluster (Cell cluster with overall highest *Ace* expression). (**D**) Data from the ChIP–seq database indicating enrichment of Fkh at the promoter of *Ace*. (**E**) JBrowse viewer of *Ace* gene with Janelia Gal4 line putative brain enhancer regions and REDfly transcriptional regulatory regions. (**F**) Relative *Ace* expression in guts of indicated genotype via RT-qPCR. n=7 replicates of 10 pooled guts from 2 independent experiments. (**G**) Representative images of decreased Fkh nuclear staining in CF model guts with *fkh* RNAi. scale bars 25µm. (**D**) Quantification of decreased Fkh staining in CF model guts with f*kh* RNAi. n=5 from 1 independent experiment. pValues were calculated using the Mann-Whitney test in GraphPad prism. Error bars are mean with 95% CI. (**I**) Representative images of decreased phospo-4EBP1 staining in CF anterior midguts compared to WT. scale bars 25µm. (**J**) Quantification of phospho-4EBP1 staining in WT and CF deficient guts. n=10 from 2 independent experiments. p values were calculated using the Mann-Whitney test in GraphPad prism. Error bars are mean with 95% CI. *Drosophila* image in panel B is from bioicons licensed under CC0.
