## Supplementary material for "Cholinergic Signaling Modulates Intestinal Pathophysiology in a *Drosophila* Model of Cystic Fibrosis": Table S1

| TF2TG | | | | | | | |
|---|---|---|---|---|---|---|---|
| **TF FBgn** | **TF Symbol** | **Peak Count** | **Motif Count** | **REDfly TFBS Count** | **Location** | **Protein-Protein Interactors** | **Genetic Interactors** |
| FBgn0000659 | fkh | 1 | 2 | 0 | intragenic | Ada2b | |
| FBgn0000659 | fkh | 1 | 2 | 0 | upstream | Ada2b | |
| FBgn0035625 | Blimp-1 | 2 | 5 | 0 | upstream | | |
| FBgn0035625 | Blimp-1 | 0 | 7 | 0 | intragenic | | |
| FBgn0011236 | ken | 0 | 1 | 0 | intragenic | pzg, Ada2b, E(bx), Trl | |
| FBgn0011236 | ken | 0 | 1 | 0 | upstream | pzg, Ada2b, E(bx), Trl | |
| FBgn0000286 | Cf2 | 0 | 1 | 0 | intragenic | bin | |
| FBgn0027339 | jim | 0 | 20 | 0 | intragenic | | |
| FBgn0027339 | jim | 0 | 1 | 0 | upstream | | |
| FBgn0283451 | br | 1 | 0 | 0 | upstream | Ada2b, Rel, rib | Met (pubmed) |
| FBgn0283451 | br | 0 | 1 | 0 | intragenic | Ada2b, Rel, rib | Met (pubmed) |
| FBgn0264490 | Eip93F | 0 | 1 | 0 | intragenic | | |
| FBgn0261113 | Xrp1 | 1 | 0 | 0 | upstream | Irbp18 | |
| FBgn0261113 | Xrp1 | 0 | 2 | 0 | intragenic | Irbp18 | |
| FBgn0085424 | nub | 0 | 5 | 0 | intragenic | pdm2, wek | |
| FBgn0085424 | nub | 0 | 4 | 0 | upstream | pdm2, wek | |
| FBgn0032223 | GATAd | 0 | 5 | 0 | upstream | mam, Ada2b | |
| FBgn0262582 | cic | 2 | 0 | 0 | upstream | gro | Dl (pubmed) |
| FBgn0262582 | cic | 0 | 1 | 0 | intragenic | gro | Dl (pubmed) |
| FBgn0005660 | Ets21C | 0 | 1 | 0 | upstream | | |
| FBgn0014018 | Rel | 2 | 1 | 0 | upstream | htk, Dif, dl, pzg, br, grh, sqz | |
| FBgn0004394 | pdm2 | 0 | 3 | 0 | intragenic | salr, nub | |
| FBgn0004394 | pdm2 | 0 | 4 | 0 | upstream | salr, nub | |
| FBgn0032587 | CG5953 | 0 | 1 | 0 | upstream | knrl | |
| FBgn0263108 | BtbVII | 0 | 1 | 0 | intragenic | CG32121 | |
| FBgn0013263 | Trl | 3 | 0 | 0 | intragenic | CG12155, CG8924, psq, Ssrp, ken, pzg, E2f1, bab2, lola, E(bx), Gug, Adf1, ttk | |
| FBgn0013263 | Trl | 2 | 0 | 0 | upstream | CG12155, CG8924, psq, Ssrp, ken, pzg, E2f1, bab2, lola, E(bx), Gug, Adf1, ttk | |
| FBgn0264490 | Eip93F | 0 | 1 | 0 | intragenic | | |

**Supplementary Table 1: Related to Figure 5**

TF2TG results for TFs with ChIP-seq Peak counts and/or binding motifs within 5 kb of Ace gene that were included in the transcription factor screen for Ace transcription (Fig 5A) .
